# Natural and age-related variation in circulating human hematopoietic stem cells

**DOI:** 10.1101/2023.11.30.569167

**Authors:** N. Furer, N. Rappoport, O. Milman, A. Lifshitz, A. Bercovich, O. Ben-Kiki, A. Danin, M. Kedmi, Z. Shipony, D. Lipson, E. Meiri, G Yanai, S. Shapira, N. Arber, S. Berdichevsky, S. Tavor, J. Tyner, S. Joshi, D. Landau, S. Ganesan, N. Dusaj, P. Chamely, N. Kaushansky, N. Chapal-Ilani, R. Shamir, A. Tanay, LI Shlush

## Abstract

Hematopoietic stem and progenitor cells (HSPCs) deliver life-long multi-lineage output. However, with aging, we exhibit certain characteristic blood count changes and accumulation of clonal disorders. Better understanding of inter-individual variation in HSPC behavior is needed to understand these age-related phenomena and the transition from health to chronic and acute hematological malignancies. Here we study 627K single circulating CD34+ HSPCs (cHSPCs) from 148 healthy individuals, along with their clinical information and clonal hematopoiesis (CH) profiles, to characterize population-wide and age-related hematopoietic variability. Individuals with CH were linked with reduced frequencies of lymphocyte progenitors and higher RDW. An age-related decrease in lymphoid progenitors was observed, predominantly in males. Inter-individual transcriptional variation in expression of a Lamin-A signature and stemness gene programs were linked with aging and presence of macrocytic anemia. Based on our model for healthy cHSPC variation we construct the normal reference for cHSPC subtype frequencies. We show how compositional and expression deviations from this normal reference can robustly identify myeloid malignancies and pre-malignant states. Together, our data and methodologies present a novel resource, shedding light on various age-related hematopoietic processes, and a comprehensive normal cHSPC reference, which can serve as a tool for diagnosing and characterizing hematological disorders.

## Introduction

The basis for understanding and defining human pathophysiological states is a detailed description of inter-individual heterogeneity among healthy individuals. Large population studies have identified wide inter-individual differences in complete blood counts (CBCs) of healthy individuals^1^ and exposed different age-related blood count changes, such as high RDW, macrocytic anemia, and a reduction in absolute lymphocyte counts^2^. The establishment of reference values, or population-wide normal ranges for certain blood parameters, has been crucial for patient evaluation, diagnosis and treatment. This is especially true in the field of hematology, where multiple parameters are measured simultaneously, and diagnosis is usually dependent upon the coordinated interpretation of more than one test^3^.

While CBC reference ranges are used in the clinic daily, the equivalent reference range for hematopoietic stem and progenitor cells (HSPCs) has not been established so far. As HSPCs mainly reside in the bone marrow (BM), access to these cells, especially in the healthy population has been problematic, whereas their general paucity in the circulation made it quite challenging to characterize them efficiently from the blood. This has become feasible given modern technologies such as single cell RNA sequencing (scRNAseq), allowing high resolution expression-based quantification and characterization of these rare cellular states.

Individual heterogeneity in the frequency of circulating HSPCs (cHSPCs) has been reported in the past, and was linked to age, smoking, sex, and hereditary factors^8^, as well as to different pathological states^9^. Few studies analyzed HSPC heterogeneity in higher resolution, but their sample size was limited^10^. Previous studies have demonstrated that most HSPC subpopulations can be identified in the PB^11^, including some based on scRNAseq analysis^12^. Functional stem cells were identified in the PB of mice^13^ and humans^12^, but, as the PB connects the BM to other extramedullary sites, it could also be enriched in unique stem cell populations^9^.

Here we apply scRNAseq analysis to cHSPCs from 148 healthy age- and gender-diverse individuals, to capture key aspects of interindividual heterogeneity. We describe transcriptional programs of 626,966 single CD34+ cells and how these correlate with various clinical attributes. This gave rise to a high-resolution model of cHSPCs, including a spectrum of states, from HLF/AVP positive hematopoietic stem cells (HSCs), through early common myeloid and lymphoid progenitor states, and more specified erythrocyte, basophil-eosinophil-mast, GMP, B, NK/T and DC progenitor populations. All data can be observed in https://apps.tanaylab.com/MCV/blood_aging/. Notably, we discovered extensive inter-individual heterogeneity in the frequency of specific cHSPC subtypes and found these to correlate with certain blood count parameters, presence of CH and aging. We further developed various tools for evaluating patient blood through compositional analysis (equivalent to the classical CBC, but at the HSPC level), followed by in-depth analysis of patient gene expression signatures, and karyotype analysis. These tools greatly enhance and streamline the analysis of hematological disorders, as we show in several case studies.

## Results

### Universal stem and progenitor states observed across humans in CD34+ peripheral blood

To evaluate interpersonal diversity in subtype distribution and regulation of cHSPCs in healthy humans, we combined multiplexed scRNAseq with genotyping, and integrated clinical data. Multiplexing was resolved using SNPs we identified in the 3’ UTR of cHSPC RNA, facilitating precise matching of cells to individuals, and improving control for batch effects and doublets (**Fig 1A**). Altogether, we collected HSPCs from 79 males and 69 females between the ages of 23 and 91 years (median 61.5) (**EDF 1A**, **Supplementary Table S1**). We ran technical replicates on 39 individuals, and biological replicates on a follow-up cohort of 20 individuals (1 year following their original sampling date). We collected longitudinal CBCs up to 5 years prior to scRNAseq along with various other clinical parameters, and performed deep targeted somatic mutation analysis on DNA produced from their blood at sampling, to identify cases of CH (**Supplementary Tables S2, S3**)^14^. Following quality control and filtering, we retained 846,762 single cell profiles, which were normalized to control for sequencing-platform batch effects and combined to construct and annotate a metacell manifold model^15^ (**EDF 1B,C**). We retained 672,000 CD34+ single cells for downstream analysis (**EDF 1D)**. These formed a rich repertoire of states, associated with cHSCs and their differentiation trajectories (**Fig 1B**, **EDF 1E,F**). The derived model recapitulated and deepened earlier characterization efforts of HSPC states from the BM, and while not fully reflecting BM dynamics, was compatible with our own BM scRNAseq data and that of others^16,17^ (**EDF 2,3A**), suggesting that cHSPCs can serve as a highly accessible proxy for hematopoietic dynamics, both within and between individuals. One notable characteristic specific to cHSPCs, however, was the repression of cell cycle gene expression (**EDF 3B**). Importantly, we found our cHSPC model to be consistent among individuals. The median number of individuals contributing cells to each metacell was 84, and all metacells included cells from at least 47 individuals. Individual-specific differential expression was limited after controlling for each sample’s cell distribution over the atlas states (**EDF 4A,B**).

**Figure 1.**
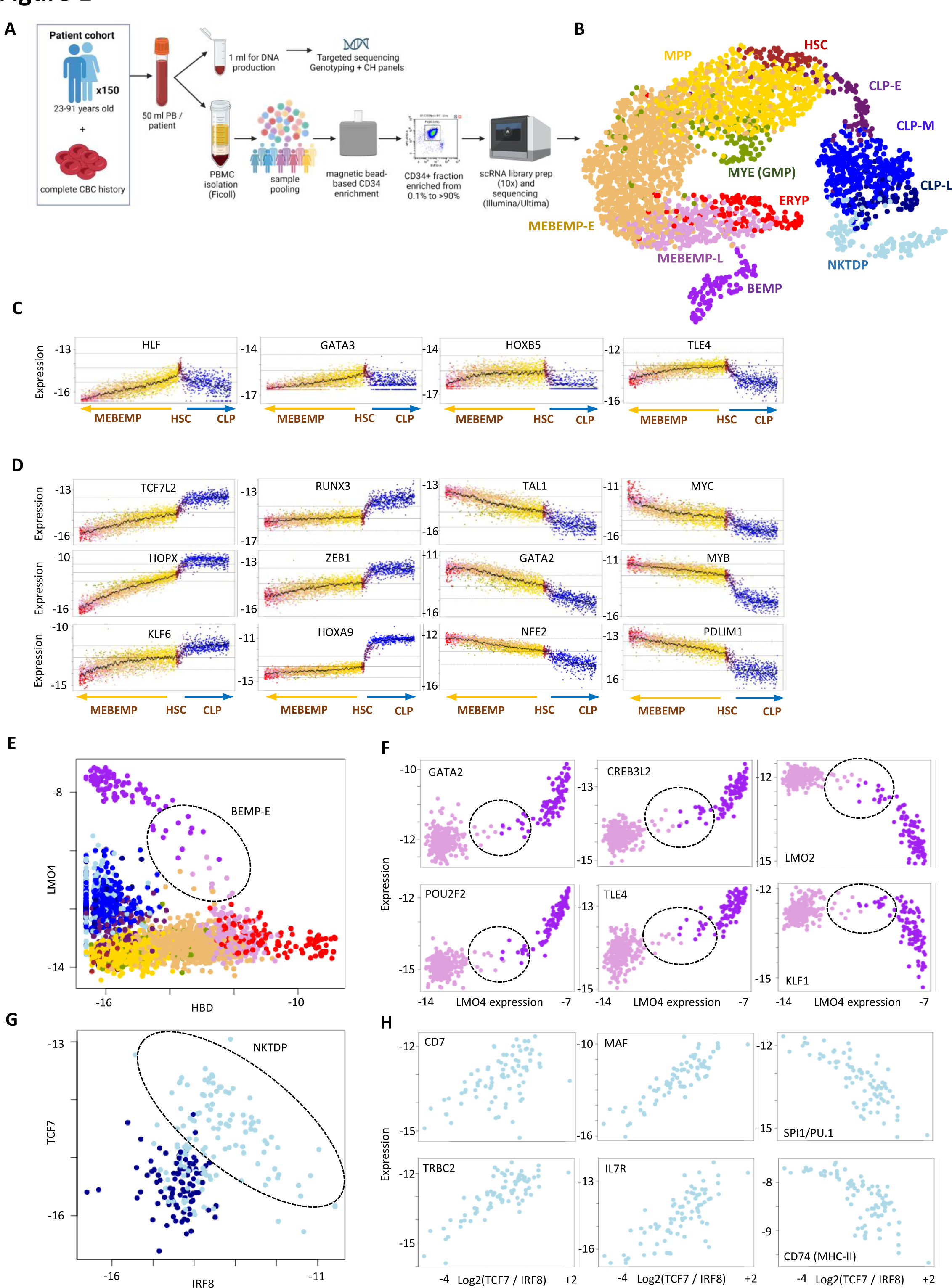
**A** – experimental design, **1B** – annotated 2D UMAP projection of our metacell manifold following filtration of metacells with low CD34 expression. Symmetric (**1C**) and asymmetric (**1D**) regulation of specific HSC transcription factors upon bifurcation to the CLP (right) and MEBEMP (left) lineages. Each panel shows the expression of one gene (y axis). Metacells in all panels are ordered (left to right) by increasing AVP expression in the MEBEMP lineage and decreasing AVP expression in the CLP lineage. Units for gene expression in all figure panels are log2 of each gene’s fractional expression. **1E** – the metacell population of interest (dotted line) linking BEMPs to their MEBEMP-L precursors. **1F** – positively and negatively regulated TFs involved in early BEMP differentiation. **1G** – gene-gene plot of *IRF8* against *TCF7* expression as hallmark markers of DC and T cell differentiation respectively. The high *ACY3* NKTDP metacell population of interest is depicted (dotted line). This population exhibits high expression of both T and dendritic cell regulators, forming a gradient consisting of NK/T cell-like progenitors exhibiting a high *TCF7/IRF8* expression ratio along with high expression of other T cell hallmarks such as *CD7*, *MAF*, *IL7R*, *TRBC2*, and DC-like progenitors exhibiting a low *TCF7/IRF8* expression ratio, along with high expression of other DC hallmarks, such as the myeloid TF *PU.1* and the MHC class II gene *CD74* (**1H**).

### High-resolution cHSC map shows *HLF*, *GATA3*, *HOXB5* and *TLE4* as distinct HSC TFs

One of the hallmarks of our cHSPC model is a distinct HSC state that is transcriptionally linked with two major differentiation gradients: The first representing a continuum of common lymphoid progenitor (CLP) programs; The second, and more common branch, representing multipotent progenitor (MPP) states and their differentiation toward granulocyte-monocyte progenitors (GMP), erythrocyte progenitors (ERYP) and basophil/eosinophil/mast progenitors (BEMP). Technical limitations of cell disassociation in scRNAseq prevented precise megakaryocyte program modeling (**EDF 4C**). We therefore annotated states at the base of this trajectory as megakaryocyte/erythrocyte/basophil/eosinophil/mast progenitors (MEBEMP) as these are also presumed to be the cells of origin of megakaryocytes^12^.

Early HSCs are marked by high *AVP* and *HLF* expression, and were previously shown to represent a rare cell population enriched with self-renewal capacity in both BM and cord blood^18^. Our model included data on ∼14,440 *HLF*/*AVP* HSCs that could be matched with cells from independent BM atlases^16,17^, suggesting that under steady-state, HSCs with potential self-renewal capacity are present in the PB (**EDF 5A**). Together with HLF and AVP, we discovered 14 genes expressed at least 1.75-fold higher in HSCs compared to their two immediate differentiation branches (**EDF 5B, Supplementary Table S4**). We specifically identified several transcription factors (TFs) enriched in HSCs, including the genes *HOXB5*, *TLE4,* and *GATA3* (**Fig 1C**). *GATA3* was previously reported to regulate self-renewal in mice long-term HSCs^19^, yet its role in human HSCs has not been studied thus far. We note that while the HSC state is defined by unique markers that are symmetrically down-regulated upon exit to the CLP and MEBEMP trajectories (**Fig 1C**), it also expresses several lineage-specific regulators at intermediate levels, which are bifurcating anti-symmetrically upon exit from the HSC state to the CLP and MEBEMP trajectories (**Fig 1D, EDF 5B**). This may suggest that the multipotent capacity of HSCs is associated with intermediate expression of multiple regulators which is resolved with differentiation.

### NK-T-dendritic and basophil-eosinophil-mast progenitors are enriched in cHSPCs

The cHSPC atlas was enriched for basophil-eosinophil-mast progenitors (BEMP), mapped as one possible terminus of HSC differentiation. While classical studies linked these cells with a granulocyte/monocyte progenitor (GMP) origin, more recent studies suggested these emerge, at least in part, from erythroid progenitors in both mice and humans^12,20^. Our analysis allowed us to focus on a small population of metacells linking BEMPs with their MEBEMP-L precursors (**Fig 1E**). This highlighted TFs (**Fig 1F**) and other factors (**EDF 6A**) positively or negatively regulated in this postulated early stage of BEMP specification. Another rare HSPC population we could now focus on included lymphoid states with high *ACY3* expression and intermediate-to-low *DNTT* levels, a combination rarely found in human BM but present in PB (**EDF 6B**). Interestingly, we observed co-variation of key T cell regulators within this population, but also anti-correlation of these factors with some hallmarks of a dendritic cell (DC) program. This can be demonstrated by comparison of *TCF7* and *IRF8* expression (**Fig 1G**), and the matching *TCF7*-coupled dynamics of *CD7*, *MAF*, and *IL7R*, or *IRF8*-coupled dynamics of the myeloid TF *SPI1* (PU.1) and multiple MHC-II genes (**Fig 1H, EDF 6C**). We therefore termed this population NK/T/DC progenitors (NKTDP)^21,22^. To summarize, our map of cHSPCs showed a rich spectrum of differentiation trajectories and progenitor states that refined previous analyses, and a remarkable universality of states, which provided an opportunity for deciphering inter-individual hematopoietic variability.

### Inter-individual variation in cHSPC stemness and in lymphoid/myeloid differentiation bias

To study inter-individual cHSPC variation, we first looked at individual-specific cell state compositions. This was performed by quantifying cell state relative frequencies within each individual’s single-cell ensemble **(Fig 2A)**. These frequencies varied extensively as shown in **Fig 2B**. For example, HSCs and CLP-Ms, representing 2.4% and 12.6% of the CD34+ population on average, respectively, showed standard deviations of 1.0% and 6.8%, respectively. The abundant MPP and MEBEMP-E states (mean frequency of 20.7% and 37.6%, respectively) showed smaller relative variations (SD 4.9% and 5.8%, respectively). To analyze the stability of cell state frequencies over time and sampling instances, we re-sampled 20 individuals one year following their original sampling date. Both lymphoid progenitor frequencies (CLP-M, CLP-L, NKTDP), and MEBEMP (MEBEMP-E, MEBEMP-L, ERYP, BEMP) frequencies were stable within the same individual across time (**Fig 2C**).

**Figure 2.**
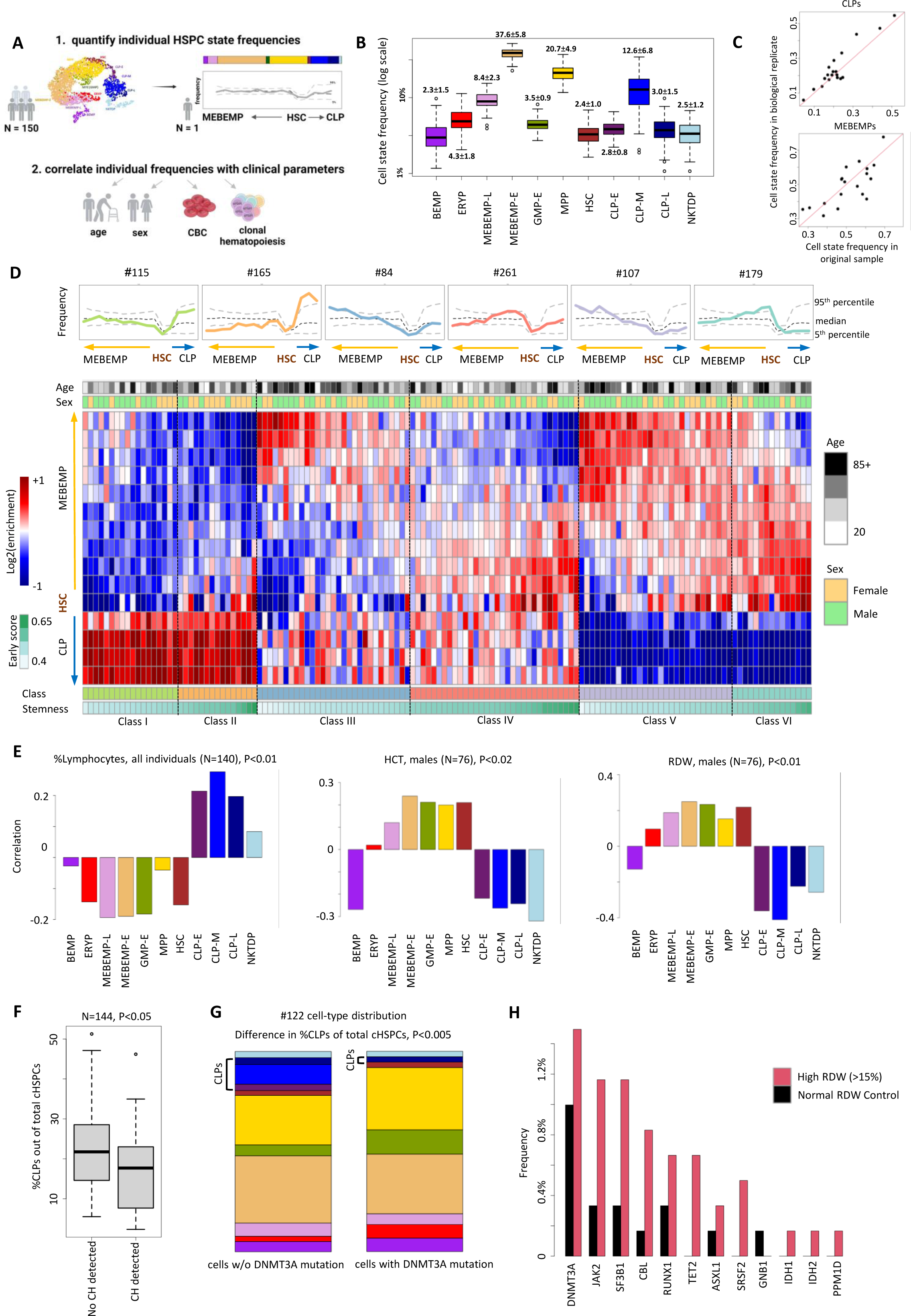
**A**- characterization of inter-individual HSPC compositional state variation (scheme). **2B** – boxplots of cell state frequency distributions across individuals (logarithmic scale). Percents calculated out of *CD34*+ population. Boxplot centers, hinges and whiskers represent median, first and third quartiles and 1.5× interquartile range, respectively. Numbers represent mean +/- SD for each distribution. **2C-** comparison of cell state frequencies between 20 biological replicates and their original samples, for CLP (CLP-E, CLP-M, CLP-L, NKTDP - top) and MEBEMP (MEBEMP-E, MEBEMP-L, ERYP, BEMP - bottom) populations. All biological replicates were sampled 1 year following original blood draw. **2D** (**top**) - individual cell state frequency profiles over the HSC-MEBEMP and HSC-CLP differentiation gradients of 6 subjects (colored lines), each representing one of six archetypes (classes) of HSPC composition in healthy individuals. Dashed lines represent the median (black) and 5^th^ and 95^th^ percentiles (grey) of the studied population. **2D** (**bottom**) cell state enrichment map over 15 differentiation bins (rows), for all studied individuals (columns) clustered into 6 classes. Classes I & II represent individuals relatively enriched in lymphoid progenitors, whereas classes V & VI represent individuals with relative depletion of lymphoid progenitors. Individuals are sorted by stemness within each class. Age and sex are denoted for each individual. **2E** – CBC correlations to cell state frequencies: %Lymphocytes (from WBC, calculated for entire cohort, left), HCT (males only, center), RDW (males only, right). Missing individuals lacked sufficient cells for analysis. Permutation test p values are displayed for each correlation. See Methods for details on the permutation-based test. **2F** – boxplots showing CLP frequency distributions in individuals with (right) and without (left) clonal hematopoiesis. **2G** – Relative cell state frequencies in mutant (right) and non-mutant (left) cells following GoT of sample #122 (DNMT3A R882 mutated, VAF = 0.07). **2H** – CH frequency (by gene) in age- and sex-matched high (red) and normal (black) RDW individuals in a cohort of 18,147 individuals.

To analyze composition in higher resolution, we profiled each individual’s enrichment over the CLP and MEBEMP trajectories. Clustering of these enrichment profiles yielded six archetypes of cHSPC composition within the healthy population (class I-VI, **Fig 2D**). These were composed of individuals with relative lymphoid enrichment (class I-II) or depletion (class V-VI) further sub-divided by a stemness gradient, enriched in classes II, IV and VI, and depleted in classes I, III and V. Analysis of technical and biological replicates confirmed this variation to be robust and patient specific (**EDF 7A,B**). To summarize, we provide the first cHSPC subpopulation normal reference range (**Fig 2B**), characterized by extensive variation among healthy individuals, and show these compositional differences are a true individual characteristic, with potential clinical implications.

### cHSPC frequencies correlate with CBCs and CH

Analysis of CBC correlations with our single-cell atlas enhanced our previous findings on the inter-individual variation in cHSPC compositions. All CBC correlation analyses were performed using median values for each blood count parameter over 5 years preceding scRNAseq. The mean and median number of blood counts per individual during this 5-year period were 8, and 6 respectively. We observed a significant positive correlation (P<0.01) between PB mature lymphocyte percentages and CLP frequencies (**Fig 2E**, left). Given the very high variation in female red blood cell counts and sizes during young adulthood (with menarche and pregnancy effects) as well as during prolonged periods of perimenopause, we analyzed RBC indices, including RBC, HCT, MCV, and RDW, separately in males and females. We observed a significant negative correlation (P<0.02) between CLP frequencies and Hematocrit (HCT, males, **Fig 2E -** middle), as well as a significant positive correlation (P<0.01) between increased RDW - a cHSPC myeloid bias - and a relative CLP depletion (males, **Fig 2E -** right). Our previous work^24^ and the work of others^25^ correlated increased RDW with high risk for CH and predisposition to AML. We demonstrate that low CLP frequencies are associated with CH (two-sided Mann-Whitney test; **Fig 2F, EDF 8A**), and further enhance this observation by performing Genotyping of Transcriptomes^26^ on one of our DNMT3A R882 cases, identifying a lower fraction of CLP cells in the mutant clone (P<0.005, Fisher’s exact test, **Fig 2G**). To further explore this association, we studied a cohort of 18,147 healthy individuals for whom we had both longitudinal CBCs and DNA available. We identified 602 individuals with a high RDW (>15%, not meeting minimal criteria for MDS) and 602 age and sex matched normal RDW controls. We performed deep targeted sequencing to identify pLMs on both high-RDW individuals and controls and found a significant enrichment of CH+ cases in the high RDW group (Fisher’s exact test P<0.002, **Fig 2H**, **Supplementary Tables S5,S6**). Altogether, the data demonstrate a 3-way linkage between decreased CLP frequencies, a high RDW, and CH.

### Age-related myeloid bias is predominantly observed in males

Blood aging is a complex and multi-factorial process, likely driven by intrinsic factors such as pre-leukemic mutations, and extrinsic effects, such as cytokine and hormonal changes. In order to decouple these factors as much as possible, we studied age-related changes in cHSPC populations in individuals without CH mutations. Analysis of age-linked compositional changes in cHSPCs within this group showed a remarkable increase in myeloid (MEBEMP) to lymphoid (CLP) ratios in males (when comparing <50 to >60-year-old individuals, **Fig 3A**). This effect was not significant in females. In this regard it is important to note that a decline in lymphocyte counts can be observed in both elderly males and females, however it appears in females at an older age^2^. Interestingly, females exhibit a surge in lymphocyte counts immediately following menopause, contributing to this delay in lymphocyte decline. Within the MEBEMP differentiation trajectory, aging was correlated with over-representation of more differentiated states, once again only in males (**Fig 3B**). Of note, the frequencies of cHSCs did not significantly change with age (**Fig 3C**). While previous studies suggested aging is linked with an increase in HSC frequencies^27^, such increase was not observed with the restrictive definitions employed here, as well as when determined from CD34+ PB HSPC frequencies in a recent cohort of 1000 healthy individuals undergoing PBMC scRNAseq^28^ (**Fig 3D**). The sex-specific correlation between age and cHSPC myeloid bias highlights the role of such non-intrinsic effects on this classical hallmark of blood aging.

**Figure 3.**
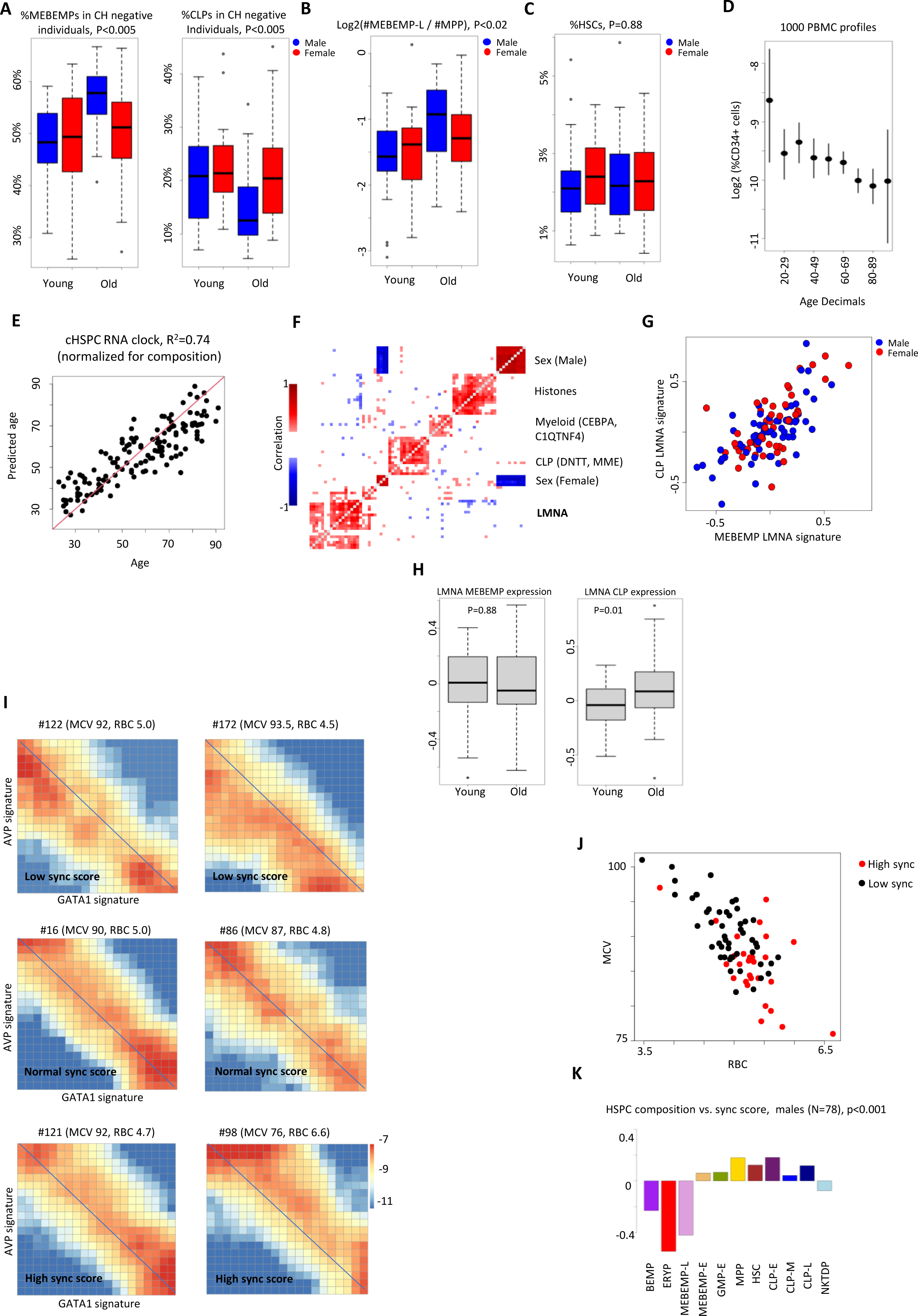
**A** – Analysis of age-linked compositional differences in MEBEMP (MEBEMP-E, MEBEMP-L, ERYP, BEMP, left) and CLP (CLP-E, CLP-M, CLP-L, right) populations, comparing specific cell state frequencies (out of total *CD34*+ population) in young (<50 years) vs. old (>60-years) individuals without clonal hematopoiesis, performed for males (blue) and females (red) separately. Kruskal-Wallis p values for group differences are denoted on top. **3B** - Analysis of age-linked compositional differences within the MEBEMP differentiation trajectory, comparing the abundance of more (MEBEMP-L) to less (MPP) differentiated states in young (<50 years) vs. old (>60-years) individuals, performed for males (blue) and females (red) separately. The Kruskal-Wallis p value for difference among these groups is denoted on top. **3C** – as in 3A, for the HSC population. **3D** – Analysis of age-linked differences in (CD34+) cHSPC frequencies from total PBMCs in a recent cohort of 1000 healthy individuals undergoing PBMC scRNAseq^28^. **3E** – True age (x) vs age predicted based on composition-controlled MEBEMP expression (y). **3F** – gene-gene correlation heatmap, calculated over individual-level HSC-MEBEMP gene expression controlled for HSC-MEBEMP composition. **3G** - intra-individual correlation of *LMNA* signatures in CLPs and MEBEMPs, for both males (blue) and females (red). **3H** – Analysis of age-linked differences in LMNA signature expression for CLP (right) and MEBEMP (left) populations in young (<50 years) vs. old (>60-years) individuals. Y axis denotes log2(observed/expected expression) normalized for composition. Boxplot centers, hinges and whiskers represent median, first and third quartiles and 1.5× interquartile range, respectively. **3I** – individual heatmaps of single cell counts over 20 bins of stemness (*AVP* signature, y axis) and MEBEMP differentiation (*GATA1* signature, x axis). Individual identifier, as well as his/her RBC, and MCV are denoted on top. **3J** – comparison between individual sync scores and clinical parameters (RBC/MCV) across males. High and low sync scores (denoted by red and black dots respectively) define clinically distinct populations. **3K** – correlation between individual sync scores and cell state compositions. Permutation test p value denoted on top.

### Composition-controlled HSPC expression correlates with age

As shown above, an individual’s cHSPC composition provides an initial blueprint of hematopoietic dynamics along the stemness and CLP/MEBEMP axes. Further analysis of transcriptional variation could now be carried out, while controlling for the dominant effect of cHSPC composition, in order to characterize additional gene expression signatures that could distinguish between individuals. Composition-controlled individual expression profiles showed high information content when correlated with age, enabling age prediction based on normalized expression alone (**Fig 3E**, **EDF 8B; see Supplementary Tables S7,S8 for additional screening for age-, CBC-, CH- and sex-associated gene expression**). We next looked for gene groups (signatures) that co-variate between individuals, filtering out sex-linked signatures and those showing strong batch effects. The most prominent of these signatures included *Lamin-A (LMNA)* as well as ANXA*1*, *AHNAK*, *MYADM, TSPAN2,* and *VIM,* among others (**Fig 3F**, **EDF 8C,D**, **Supplementary Table S**9). Individual LMNA signature expression varied across a range of more than 2-fold (**EDF 8E**), exhibiting high expression variability in HSCs and early myeloid and lymphoid cell states, and a homogeneously low expression in late MEBEMPs and CLPs (**EDF 8F**). Individual LMNA signature expression was consistent in myeloid and lymphoid cell states (**Fig 3G**) and was stable in our follow-up cohort (**EDF 8G**). Interestingly, we observed an age-linked increase in LMNA signature expression in lymphoid, but not myeloid, cHSPCs (**Fig 3H**). Taken together, we show that in addition to the accumulation of pLMs in HSPCs, aging is strongly linked with changes in the distribution of progenitor cell states in the PB, and with significant differences in the expression of certain gene signatures. The mechanistic basis for this variation and its clinical impact remain unresolved.

### Rapid repression of stemness signatures in MEBEMPs is linked with lower red cell counts and higher red cell volumes

The differentiation of HSPCs toward MEBEMP and CLP fates involves coordinated activation and repression of specific transcriptional programs that were generally universal among individuals. Yet, our screen for inter-individual variation in gene signatures suggested that individuals differed in the way they synchronized the opposing effects of these stemness and differentiation programs. To quantify this variation, we compared *AVP* (stemness) and *GATA1* (MEBEMP differentiation) signatures (**Supplementary Table S10**) on a 20×20 bin expression matrix (**Fig 3I**). While most individuals displayed dynamics close to the diagonal line (individuals #16, #86, for example), following the typical transition from stemness to differentiation, some individuals deviated from the diagonal, indicating skewed synchronization between the *AVP* and *GATA1* signatures. We quantified this deviation (i.e. off-diagonal frequency) using a new synchronization-score. This facilitated the identification of individuals with sync-scores as low as 0.12 (#122 and #172, for example, **Fig 3I**, top), indicating delayed activation of *GATA1* relative to *AVP* repression. Namely, while these individuals rapidly reduce their *AVP* expression, their increase in *GATA1* and *GATA1*-related genes is delayed. In contrast, individuals exhibiting a high sync-score (#98 and #121, for example, **Fig 3I**, bottom), show early activation of *GATA1* expression, which precedes *AVP* repression. We detected stability of the sync-score in our follow-up cohort (**EDF 8H**). Inter-individual sync score variability was positively correlated with RBC levels, and consistently anti-correlated with MCV in males (P<0.01 (spearman) for both RBC and MCV, **Fig 3J**). Analysis of the correlation between individual sync-scores and cHSPC compositions in males demonstrated a negative correlation with ERYPs and BEMPs (**Fig 3K**). To summarize, we demonstrated variation in the coordination of stemness and MEBEMP differentiation programs that is correlated with red blood cell counts and volumes. Hypotheses on the mechanisms underlying this heterogeneity and its greater clinical consequences should be further explored. Individual scores and signatures (LMNA signature in MEBEMPs and CLPs, sync-scores, and HSPC cell state distributions) are summarized in **Supplementary Table S11**.

### Using the cHSPCs atlas for mapping, dissecting and annotating myeloid malignancies

Diagnosis of myeloid malignancies requires the identification of clonal markers (mutations / structural variants) and the detection and quantification of blasts by microscopy and flow cytometry. In **Fig 4A** we describe a new stepwise approach for analysis of myeloid disorders based on sampling of cHSPCs and comparison of their compositions, normalized expression and copy number variations (karyotyping) to our new normal reference. As proof of concept, we analyzed sampled cells from 2 new healthy individuals, 3 CMML patients, 2 MDS patients, 1 MDS/MPN overlap patient and 1 myelofibrosis patient. We further sampled and analyzed 2 AML cases to demonstrate how acute disease is manifested when projected over the cHSPC model (**EDF 9-11**). Projection of patient metacells showed a high gene expression correlation between metacell pairs (**Fig 4B**, color coded dots), for all pathological cases except for the AMLs. Cell state compositions were, however, skewed in all pathological cases (**Fig 4C,D**), but not for the healthy controls. All patient samples showed a remarkable reduction in CLP populations (**Fig 4C,D**). Two of the CMML patients (N192, N235) demonstrated highly abnormal enrichment of specific (basophil and myeloid) cell states, while the rest showed a relatively balanced distribution over the MEBEMP differentiation spectrum (with slight enrichment of stem states). Compositional-controlled gene expression comparisons of patient samples to the normal model identified specific genes that were recurrently induced or repressed in disease (**Fig 4E**). While both healthy individuals and the treated MDS case showed a minimal number of DE genes, all leukemic cases exhibited a substantial increase in the number of DE genes when compared to the healthy population model (**Fig 4E**).

**Figure 4.**
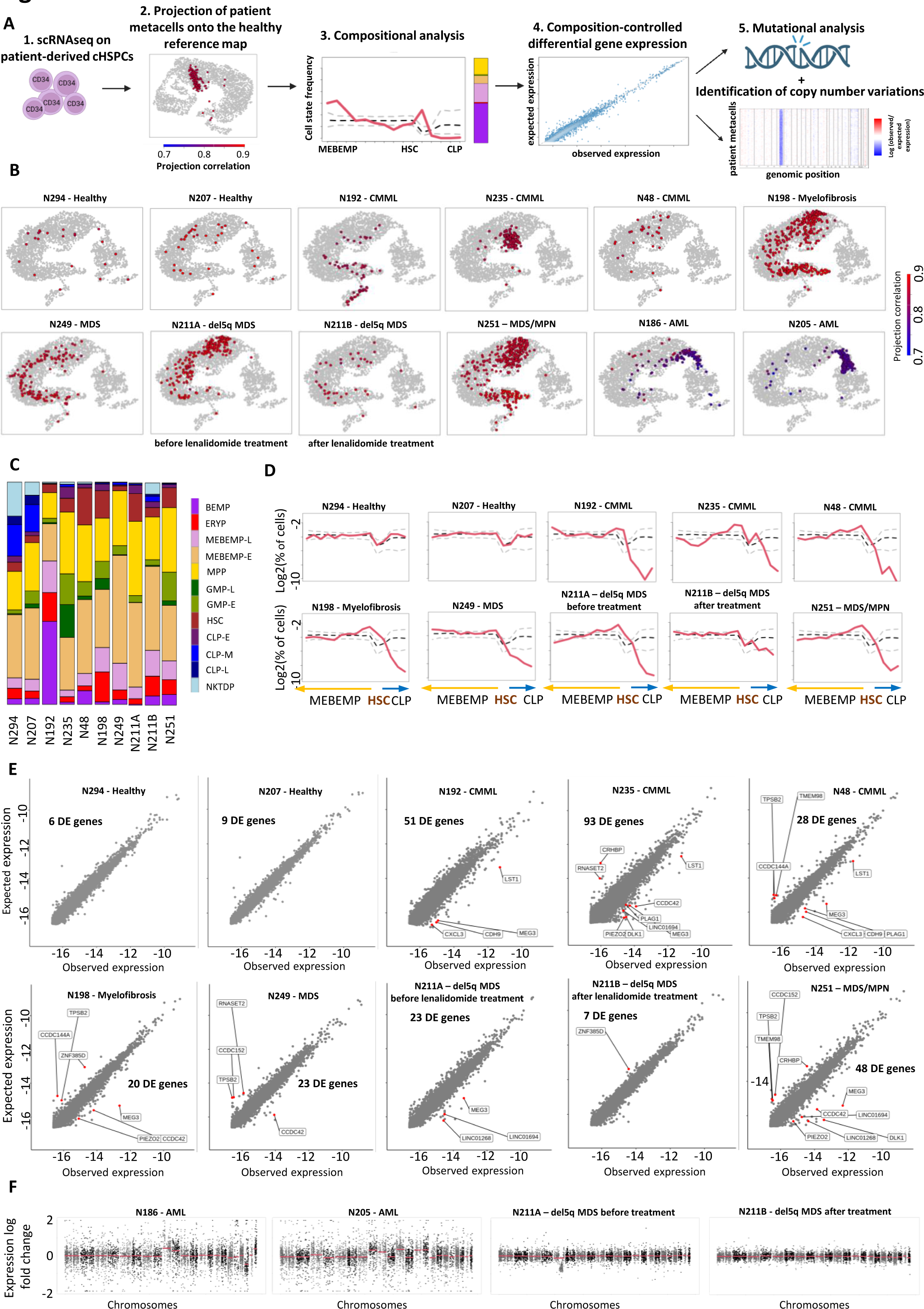
**A** – diagnostic approach to leukemia analysis using our cHSPC reference atlas (scheme): 1. scRNAseq on CD34-enriched PB and construction of a patient-specific metacell model, 2. Projection of patient derived metacells on the healthy reference atlas. 3. Compositional (relative cell state frequency) analysis and 4. Composition-controlled differential gene expression analysis. 5. Mutational and CNV analysis using targeted DNA sequencing and RNA-based karyotyping. **4B** –projection of metacells derived from 2 new healthy individuals (not included in the reference model), 3 CMML patients, 2 MDS patients (1 of them prior to and following treatment), 1 MDS/MPN overlap patient, 1 myelofibrosis patient and 2 AML patients on our healthy HSPC reference metacell model. Gene expression correlations between patient (projected) and reference metacells are color coded according to the legend on the right. **4C,D** – individual cell state frequency profiles over the HSC-MEBEMP and HSC-CLP differentiation gradients for 2 healthy and 7 patient samples (1 of them prior to and following treatment initiation). Dashed lines in Figure 4D represent the median (black) and 5^th^ and 95^th^ percentiles (grey) of the healthy population. **4E** – Compositional-controlled differential gene expression of 2 healthy and 7 patient samples (1 of them prior to and following treatment initiation) to the normal reference, quantifying the number of differentially expressed genes and identifying specific genes recurrently induced or repressed in disease. **4F** - scRNAseq karyotyping for 2 AML patients and 1 MDS patient prior to and following treatment initiation. Metacell models were created for each MDS/AML patient and projected over our healthy reference map. Coupled reference and projected (patient) metacells were then used for calculating expression ratios over all expressed genes in all chromosomes. Log2 fold-change expression (patient/healthy) for all expressed genes across all chromosomes is shown. Red lines represent the median of each chromosomal fold-change distribution.

Detection of karyotypic abnormalities based on gene expression dosage effects, previously suggested and implemented in several tools^29^, can be readily implemented on cHSPCs, as shown in **Fig 4F** and **EDF 9A**,**B**, where clear chromosomal aberrations are observed in the two AML cases analyzed. We further detected deletion of the long arm of chromosome 5, del(5q) in one of our MDS samples with complete cytogenic remission following lenalidomide treatment (N211A & N211B, **Fig 4F**, **EDF 9A**). To conclude, the atlas of normal cHSPC states presented herein enables high resolution characterization of myeloid disorders, based on their cell states, compositional-controlled transcriptional variance, and abnormal karyotypes.

## Discussion

This study characterizes interindividual heterogeneity in circulating HSPCs across 148 healthy individuals, analyzing 627K PB CD34+ cells via scRNAseq. The magnitude of our cohort, along with the potency and resolution of modern single cell technologies and the computational methods used here, allowed us to characterize in detail the transcriptional programs of diverse, sometimes rare (NKTDP, BEMP), HSPC sub-populations, refining and augmenting previous findings from much smaller cohorts (**Fig 1**). We define a normal reference range for cHSPC subpopulation frequencies within a large age- and sex-diverse healthy population and show that cHSPC subtype compositions are highly variable between individuals, while the cell states themselves are remarkably universal (**Fig 2**). These compositions remained stable over a one-year follow-up period, placing them as a strong individual characteristic. Future studies will need to further explore and better define the mechanistic and genetic basis for this compositional heterogeneity. At the population level we discovered a significant correlation between low CLP frequencies, CH and increased RDW (**Fig 2**). We further show that the known age-related myeloid bias in HSPCs is significantly male driven (**Fig 3**). Analysis of composition-controlled transcriptional variance identified an RNA expression clock which correlated with age (**Fig 3**), and a novel gene module (*LMNA*) whose expression increased in CLPs with age. We discovered that healthy individuals regulate the transition from stemness to myeloid differentiation differently and found that premature down-regulation of stemness genes correlates with high MCV anemia (**Fig 3**). We conclude by demonstrating how this resource can be used to effectively identify and diagnose pathological cases based on their abnormal cHSPC subpopulation frequencies, differential gene signatures, and chromosomal aberrations (**Fig 4**), all from PB samples.

The nature and magnitude of our cohort allowed us to produce **Fig 2B**, which in our opinion, presents the key finding of this study - the first circulating HSPC subtype normal reference range in a healthy population, which is both age and sex diversified. Though the relation between circulating HSPCs and those found in the BM has not been completely characterized yet, our PB-BM comparisons provide evidence for the consistency of our model. Our data demonstrates that cHSPCs resemble their BM counterparts transcriptionally, with the exception of lower cell cycle gene expression (**EDF 2,3**). Importantly, we are *not* arguing that cHSPCs are (or should be) a complete model for hematopoiesis in the BM, but that they are nonetheless useful proxies for major processes driving hematological physiology and pathology – and in this context - their accessibility and robustness are of major utility. Inter-individual differences in cHSPC compositions and states can serve as an indispensable mean for capturing key aspects of a patient’s hematopoietic state. The relevance and importance of this normal reference can perhaps be better understood in view of the normal CBC reference range, as only once a population-wide reference was set for CBCs, could deviations from it be considered pathological. In **Fig 4** we provide a detailed pipeline for the identification and characterization of such pathologies based on our normal reference. We fully acknowledge that our reference model is not universal, that it should be explored in diverse populations and in a wider range of pathologies, and that as a resource it surely can and will be further improved.

To conclude, our study delves into the basic molecular physiology of cHSPCs at the population level, uncovering age-related phenotypes and proposing a platform for novel mechanistic and diagnostic insights into blood malignancies. This, along with various other novel tools for profiling genetics and epigenomics in the blood, will redefine normal vs. pathological states in hematology and provide both clinicians and researchers new means for mapping the transition from health to disease.

## Methods

### Sample procurement and handling

We collected fresh peripheral blood samples from 148 healthy individuals (79 males, 69 females) aged 23-91. Their demographics and molecular data are presented in **Supplementary Tables S1 and S3** (Basic participant demographics and CH status). All sample donors were considered healthy, their CBCs were within normal range, and they were not known to have any CH defining mutations prior to sequencing. Written informed consent allowing access to longitudinal CBCs and sequencing data (CH and genotyping panels) was obtained from all participants in accordance with the Declaration of Helsinki. All relevant ethical regulations were followed, and all protocols were approved by the Weizmann Institute of Science ethics committee (under IRB protocol 283-1).

Recruitment was intended to allow characterization of the normal variation in circulating HSPC states. As no such profiling had been previously performed, we could not assume much regarding the variance in the population a priori. We therefore aimed to profile responsive volunteers with normal blood counts, balancing sex and seeking a dispersed age distribution biased toward older individuals. We reassessed this strategy following initial sampling, and observed remarkable homogeneity in transcriptional states across individuals sampled from our immediate community as well as from HMO out-patient clinics, emergency medical centers, hospital wards etc. This was critically important, as it confirmed the universality of our model across individuals. We observed large variation in the composition of states across healthy individuals, which suggested that more sampling, without specific preferences, would be appropriate for further characterizing compositional variation. Our results support two characteristics of human cHSPCs that we believe to be nontrivial and robust for alteration in sampling strategies: the universality of molecular cell states among individuals (allowing one to use the model as a reference for any patient), and the high degree of variation in composition between healthy individuals (allowing one to look at interesting clinical populations vis-à-vis our model for normal variation).

50 ml of PB were drawn from each individual into lithium-heparin tubes. 1 ml of blood was used for DNA production, and the remaining volume was used for PBMC isolation via Ficoll, using Lymphoprep filled Sepmate tubes (StemCell technologies), followed by CD34 magnetic bead-based enrichment using the EasySep human CD34 positive selection kit II (StemCell technologies). We found this enrichment strategy to be simple and reproducible and chose it for several reasons: 1) RNAseq data was most reproducible when cells were not sorted, but rather enriched-for using beads (lower mitochondrial gene fraction). 2) CD34 purity could be highly regulated by this method, to achieve anywhere between 50-95% enrichment of CD34-positive cells, which could later be easily distinguished based on their single cell expression data. In terms of cell numbers - 50 ml of blood would yield anywhere between 50 to 100 million PBMCs following Ficoll, 1/1000 of which are expected to be CD34+, such that we increased this population’s representation from 0.1% in the periphery to at least 50% of cells loaded for analysis.

### ScRNAseq of CD34+ PBMCs

Single cell RNA libraries were generated using the 10x genomics scRNAseq platform (Chromium Next Gem single cell 3’ reagent kit V3.1). Chip loading was preceded by flow-cytometry to verify that enrichment was successful, and that enough CD34+CD45^int^ live cells were gathered. All blood samples were freshly drawn at the Weizmann Institute of Science on the morning of each experiment day, and time from blood draw to 10x loading was restricted to 5 hours. The motivation for working with fresh samples was based on our previous experience with PB CD34+ cells being vulnerable to freezing/thawing rounds and long manipulation times.

10x libraries were sequenced on two alternative platforms (Illumina / Ultima Genomics). 12 libraries were simultaneously sequenced on both platforms for comparison purposes and in order to demonstrate the scalability of our approach. We observed the Ultima-sequenced data to be highly similar to the Illumina-sequenced data (**EDF 7A**).

### Genotype-based demultiplexing

All cells were traced back to their sample of origin using genotype-based de-multiplexing. This method allowed pooling of blood samples immediately following extraction of the DNA aliquot, such that CD34-enrichment was performed on the entire pool of PBMCs produced. The use of SNP-based multiplexing has several advantages to alternative antibody-based cell hashing methods: 1) it is extremely cost effective, such that the cost of sequencing a single individual on a 2000 SNP Molecular Inversion Probe (MIP) panel at a depth of 1000X per SNP (adequate for de-multiplexing purposes) is several folds cheaper than antibody staining, 2) genotyping eliminates the need to keep samples separated prior to loading, it entails shorter handling times and less cell manipulation, as it does not require antibody incubation periods and multiple wash centrifuges. This was very evident in cell viability prior to chip loading. As with other methods of sample multiplexing, genotype-based multiplexing allows for robust doublet detection during data analysis, which enabled loading of 30-40K cells from between 4-6 individuals on each Chromium Chip lane, yielding 15-25k cells per library.

### Molecular Inversion Probe (MIP) panels

Both our CH and genotyping panels are Molecular inversion probes (MIP)-based panels described in detail previously^14^. Our CH panel contains 705 probes, covering pre-leukemic SNVs and Indels in 47 genes, and is complemented by 2 amplicon sequencing reactions to cover GC rich regions in *SRSF2* and *ASXL1*. Our genotyping panel allows for the simultaneous detection of >2000 common genetic variants, all of which are extensively covered in all cell types in our data. It includes heterozygous sites with at least 5% minor allele frequency from the 1K genomes project, which were highly covered by RNA molecules in our data (at least 80 UMIs across all cells in a test 10x library), excluding sites in repetitive elements and in sex chromosomes. Both panels were designed using MIPgen^30^ to ensure capture uniformity and specificity.

### Variant calling and identification of ARCH mutations

As MIP sequencing is cost-effective yet noisy, we developed an in-house variant-calling method to identify low VAF CH events^14^.

### ARCH sequencing of high RDW samples and controls

In order to compare propensity for CH and high risk CH mutations^24^ in high RDW cases and normal RDW controls, we performed deep targeted sequencing of DNA samples from 602 high RDW (>15%) individuals, who did not show signs of anemia and whose blood count did not meet MDS criteria (11.5g/dL≤Hg≤15.5g/dL [F], 13g/dL ≤Hg≤17g/dL [M], 80fL≤MCV≤96fL, PLT≥100×10^9^/L, Abs Neut≥1.8×10^9^/L), and 602 normal RDW (11.5g/dL ≤Hg≤15.5g/dL [F], 13g/dL≤Hg≤17g/dL[M], 80fL≤MCV≤96fL, PLT≥100×10^9^/L, Abs Neut≥1.8×10^9^/L), age and gender-matched controls. Case-Control matching was performed using the R MatchIt package, balanced on age and gender, method = “nearest”, ratio = 1, from a total of 18,147 individuals with longitudinal blood counts and available DNA. All DNA samples and corresponding blood counts were received de-identified from the Tel Aviv Sourasky Medical Center (TASMC) Integrative Cancer Prevention Clinic. All DNA samples were collected after obtaining written informed consent and in accordance with the Declaration of Helsinki. All relevant ethical regulations were followed, and all protocols were approved by the TASMC ethics committee (under IRB protocol 02-130). CBCs and sequencing results of all cases and controls are presented in **Supplementary Tables S5, S6.**

### scRNAseq processing

We processed fastq files by executing cellranger with an hg-38 reference genome. We filtered cells with at least 20% mitochondrial expression and ≤ 500 UMIs from unfiltered genes.

### Doublet calling

We performed several steps to assign cells to their individuals and to detect doublets. The pipeline is made of the following steps:

1. Demultiplexing cells and calling doublets based on SNPs found in the scRNAseq data
2. Building a metacell model using cells from all the libraries, including cells previously marked as doublets, and identifying metacells made of doublet cells.
3. Identifying metacells with doublet cells based on expression of marker genes.
4. Building the final metacell model and marking metacells as doublets based on expression markers.

In the first step, we identify doublets and assign cells to individuals using Vireo and Souporcell, which cluster cells based on SNPs found in sequenced RNA molecules. We executed Vireo^26^ (preceded by running cellsnp) and Souporcell^27^ on each library separately. Both methods used SNPs from our genotyping panel^15^ which were covered by at least 20 UMIs in the library (in Souporcell – at least 10 from the major and minor allele each). We observed high agreement in doublet calling between the two methods.

In the next step, we built a metacell model with cells from all libraries, including all biological and technical replicates. This model included cells that we identified as doublets, and also included cells from individuals that are not part of this study, but were multiplexed with our libraries. The model was built with metacell2, with a target metacell size of 200 cells. We then marked all metacells where at least 35% of the cells were already marked as doublets, and all metacells that expressed unique markers of distinct cell types, as doublet metacells. All cells that belonged to a doublet metacell were then marked as doublets.

We then built an additional metacell model (see below), without the cells that were marked as doublets, and only with cells of individuals belonging to this study. In this model we further filtered a small number of metacells that expressed key marker of distinct cell types as doublets.

### Assignment of cells to individuals

Vireo^26^ and Souporcell^27^ both cluster cells based on SNPs found in the sequenced RNA, such that cells in the same cluster belong to the same individual. We observed very high agreement between the two methods in their assignment of cells into individuals. In two 10x libraries where the two methods did not agree (due to individuals with a very small number of cells), we reran the methods on a subset of the cells and a smaller target number of clusters. We mostly used Vireo’s clustering, but in some libraries we also assigned cells that were unassigned by Vireo but were assigned to an individual by souporcell, if they matched well to the genotype data (see below). For one library we took Souporcell’s assignment, because of better matching to the genotype data.

In the next step we assigned clusters of cells to the individual they originated from. To this end, we correlated the genotypes of each cell cluster, as inferred by Vireo, to all genotypes we measured using the MIP panel (using sites with sufficient sequencing depth in the MIP panel). As a control, we performed matching against the MIP genotypes of all individuals in the cohort, and not just individuals from one library. We observed in all cases a clear matching to a single individual from the expected library. The assignment also correctly identified related individuals, and the sex of the matched individual was confirmed by expression of XIST in the RNA data.

### Removal of droplets with Megakaryocyte signatures

We identified droplets with complete or partial megakaryocyte expression (at least 5% of the UMIs coming from a megakaryocyte gene program including PF4, PPBP and 131 additional genes) due to their overall high doublet rate. We then built a metacell model from the retained cells (not marked as doublets, confidently assigned to an individual, did not show megakaryocyte expression).

### Correcting for sequencing platform bias

Some of our 10x libraries were sequenced by an Ultima sequencer, and as most libraries were processed through a standard Illumina pipeline, we wished to minimize batch effects related to small variation in sequencing platform. To that end we used libraries that were re-sequenced in both platforms. We calculated an Illumina-Ultima correction factor per gene as the mean log2-fold change in expression of the gene across re-sequenced libraries. We then normalized each Ultima-sequenced library by downsampling genes with at least 0.28 log2-fold Ultima overexpression, and resampling genes with at least 0.2 Illumina overexpression. The downsampling and resampling were performed for each gene independently, across all cells in a library. The thresholds for downsampling and resampling were chosen such that the overall number of UMIs per cells remained similar. 87 genes with at least 4-fold change between Ultima and Illumina were excluded from further processing.

### Computing the reference metacell model

The model was built with metacell2, with a target metacell size of 200 cells. We marked forbidden genes such as histone genes, cell cycle related genes, ribosomal genes, stress response genes (including e.g., FOS, JUN), sex-linked genes, genes with high or inconsistent differences between Illumina- and Ultima-sequenced technical replicates, and other genes that we found to have high technical variation. These genes were not used by metacell2 when calculating gene-gene similarities, but were included in downstream analysis. We annotated the metacells using known markers as illustrated in **EDF 1C**. We excluded from most downstream analyses metacells from cell types with low CD34 expression (monocytes, B cells, T cells, NK cells, DCs, endothelial cells), and metacells which expressed marker genes from distinct cell types that we marked as doublets.

For visualization of the metacell manifold we used UMAP projection of the metacell expressioin vector over genes with specific enrichment over cell types. The cell types considered were either all cell types (for **EDF 1B**), or CD34+ cell types (for **Fig 1B**).

### BM comparison and projections

We used three BM datasets: a dataset including two CD34-enriched bone marrow cells that we collected for this study, the Human Cell Atlas (HCA) dataset^19^, and a CD34+ bead-enriched BM dataset from^30^.

We built a metacell model for the two CD34-enriched BM samples we collected using metacell2. Since these samples only collected one sample per library, we did not run Vireo and Souporcell. We marked metacells expressing marker genes of distinct populations as doublets, and excluded them from further analysis.

We have previously processed and annotated the HCA dataset in a metacell model. We downloaded the Setty et al. sequencing data and processed it by running cellranger and created a metacell model. To project our PB data, our BM data, and the Setty dataset on the HCA dataset, we correlated between the HCA metacells and the projected metacells over genes showing high variance in the HCA metacell model. We annotated each Setty metacell using the mode of the 5 most correlated HCA metacells. We annotated our BM data using the mode of the 5 most correlated HCA metacells, and using expression of gene markers. We projected metacells (from each respective model) on the HCA UMAP using the mean x and y values of the 5 most correlated HCA metacells. To compare S-phase genes between the PB and BM (**EDF 3B**), we calculated for each PB and HCA metacell its S-phase signature (mean expression of six cell cycle genes: TYMS, H2AFZ, PCNA, MCM4, HELLS, MKI67), and plot the distribution of this score across metacells for each cell type.

### HSC differentiation gene programs

To visualize transcriptional dynamics in HSC cells, we sorted MEBEMP and CLP metacells based on their AVP expression. To calculate differential expression between HSC and neighboring cell types (**EDF 5B**), we calculated the geometric mean of each gene across HSCs, CLP-M and MEBEMP-E metacells, and took the difference between HSC and MEBEMP-E, and between HSC and CLP-M.

### Differential expression between individuals unexplained by the metacell model

To create **EDF 4A,B** we compared each individual’s pooled expression profile to a matched expression profile based on the individual’s distribution across metacells. We performed the analysis separately for MPP / MEBEMPs (BEMP, EP, MEBEMP-E/L, GMP-E and MPP) and CLPs (CLP-E/M/L, NKTDP). In each group of cell types, we downsampled each cell to have 500 UMIs and summed the UMIs across all cells of each individual, normalized the sum to 1 and calculated log2, to obtain the observed expression. To compute matched expression, we downsampled each metacell to have 90K UMIs and summed all UMIs of the metacell each cell belongs to for each individual. We normalized this matched expression to sum to 1, and took log2. For **EDF 4A,B** we plotted all genes that were expressed in either the observed or matched expression in at least one individual (log2 expression > 2^-14.5 for MPP / MEBEMPs, > 2^-13.5 for the CLPs which had less UMIs), and that had at least 2-fold change between observed and matched in at least one individual. We excluded genes that exhibited strong batch effects.

### HSPC compositional analysis

To explore variance in cell type composition between individuals, we first calculated the distribution of each individual’s cells across the CD34+ cell types. To perform compositional analysis at higher resolution than cell types, we partitioned cells from CD34+ cell types into finer grained bins. We used one HSC bin, four CLP bins, and ten MEBEMP / MPP bins, for a total of 15 bins. We assigned HSC cells to bin 0, CLP-E cells to CLP bin 1, and CLP-M/L cells to CLP bins 2-4 based on decreasing AVP expression of their metacells, such that bins 2-4 had the same number of cells. We similarly assigned MPP and MEBEMP-E/L cells into 10 bins based on AVP such that these bins had an equal number of cells.

For **Fig 2D** bottom panel, we calculated the enrichment of each individual’s cells in each bin (log2 of the ratio compared to the median across individuals). We partitioned individuals into three group with different CLP numbers based on each individual’s mean enrichment across CLP bins 2-4 – those with mean enrichment > 0.5 are high CLP, those with < −0.5 are low, and the rest are intermediate. We next defined the stemness score as the ratio between the number of cells in MPP / MEBEMP bins 1-5 and the total MPP / MEBEMP number (cells in bins 1-10). Individuals with stemness score > 0.5 had enriched stemness. The combinations of CLP enrichment and stemness define the six classes shown in the figure. For visualization we further sorted individuals within each cluster based on their stemness score.

In some cases, cells from one blood sample were divided to two different 10x libraries. In this case cells from both libraries were considered together to represent the individual. Biological replicates were only considered when patients were sampled on two different days. For technical replicates, we only considered cells that appeared in a specific 10x library, even if an individual had other cells in another library from the same day. This policy holds for all analyses related to biological and technical replicates in this study.

### Test for association between cell type distribution and a numerical label

We used permutation tests to test the relation between cell type distribution and a label (age, CBC, sync-score or *LMNA* signature). We sorted 11 CD34+ cell types from late MEBEMP differentiation through HSC and to late CLP differentiation (cell types are displayed by this order in **Fig 2B**). We looked at triplets of adjacent cell types in this ordering, and calculated for each triplet the total frequency each individual has from these cell types, obtaining a vector of length 9 per individual. We then correlated each of these 9 sums to the label, and took the maximal absolute value from all these correlation values as a test statistic. We repeated this process after permuting the label 1000 or 10000 times, and used the test statistics from the permutations to derive a p-value.

### Variably expressed gene modules

We detected genes modules with high variance across individuals while controlling as much as possible for compositional variation. This involved the following procedure, applied separately for myeloid and lymphoid states (described first for myeloids):

A. we calculated for each individual the 5^th^ percentile of the number of UMIs across their cells, and downsampled the cells to this number. We then pooled all downsampled cells for each individual from the MPP metacells, normalized to sum to 1 and took log2. This gave us the observed expression profile of each individual.
B. We created an expected expression profile for each individual as follows. We partitioned the MPP metacells into 30 bins based on their AVP expression, and downsampled the metacells to 90K UMIs. For each of the 30 bins, we averaged the expression of all genes across downsampled metacells in the bin. This defined an expression profile for each of the 30 bins. To obtain an individual’s expected expression, we calculated a weighted mean of the 30 bins’ expression profiles, where the weight of each bin is proportional to the fraction of the individual’s cells from that bin, normalizing to sum to 1 and taking log2. We then calculated the difference between the observed and expected expression profiles.
C. Our data show some batch effect distinguishing samples collected in two calendric periods. As this effect could introduce co-variation between genes across individuals, we applied a correction controlling for it. This was done using a linear model fitting each gene to the patient sample collection period, its age and sex. We then subtracted the inferred period factor from the samples that were collected in the second period. We found that this approach significantly reduced emergence of gene clusters linked with calendric sample collection bias.
D. We screened for genes with high variance that were unlikely to be affected residually by the main manifold differentiation process. We removed genes with high batch effects, genes with high AVP correlation (absolute value Pearson correlation > 0.65), and genes highly correlated (absolute value Pearson correlation > 0.5) with a module of genes that were differential between the first and second library collection periods. We then calculated each gene’s variance in the difference between the observed and expected expression across individuals. As some of the variance can be explained due to sampling noise, we plotted each gene’s variance across individuals compared to its mean expression across individuals. We sorted genes by this expression value and subtracted from the variance of each gene a rolling mean of the variances of 100 neighboring genes in that ordering. We chose genes with variance at least 0.08 higher than the rolling mean variance.
E. We calculated a gene-gene Spearman correlation matrix for high variance genes, and clustered the correlation profiles using hierarchical clustering. We removed genes with low mean correlation (< 0.2) to their cluster’s genes, and then removed gene clusters with low mean correlation between their genes (≤0.25 mean correlation of all gene pairs). We also computed the gene-gene correlations while using only samples from our first library collecting period, and required gene clusters to have high mean correlation (> 0.25) between their genes using only these early samples. We removed additional gene modules that we observed were due to batch effect, or the represented traces of MEBEMP differentiation that were not normalized by approach. This resulted in **Fig 3F**.

We performed a similar (steps A-E) analysis for CLPs (**EDF 8C**), with a few differences. The analysis included cells from CLP-M metacells. The cells were partitioned into 6 bins, and the partitioning was based on the average of their DNTT and VPREB1 expression. Genes with high absolute correlation to the average of DNTT and VPREB1 were excluded. After clustering the gene-gene correlation profiles and removing genes as described for the MEBEMPs, gene clusters with remaining correlation to CLP differentiation were removed.

### Age regression

We developed age regression models for MEBEMP and CLP expression separately. To predict age, we used the difference between an individual’s observed and expected gene expression as described above. We only used genes whose minimal expression across individuals is at least 2^-14.5 for MEBEMPs and 2^-15.5 for CLPs. We trained a LASSO model using nested leave-out-sample-out cross validation: For each sample we left out, we performed a leave-one-sample-out cross validation on the remaining samples to select LASSO’s λparameter, trained a model using the selected λand made a prediction on the left-out sample.

### LMNA signature

We used the difference between an individual’s observed and expected gene expression, and correlated all genes to LMNA separately in the MEBEMP expression and CLP expression matrices. We then summed the MEBEMP and CLP correlation values, and kept genes whose sum was > 0.7. We further removed genes that exhibited high technical variance, and were left with 17 genes in the signature. To calculate each individual’s LMNA signature, we calculated for each individual the average value of the 17 genes in the observed – expected matrix, for MEBEMPs and CLPs separately.

To plot **EDF 8E**, we calculated the geometric mean of the LMNA signature genes for each individual separately in each one of the 10 MPP / MEBEMP bins described earlier for **Fig 2D**.

### GoT Analysis

GoT^24^ performed on sample #122 allowed us to mark this individual’s cells as wild-type or mutated. Due to the low VAF of #122’s DNMT3A mutation, and in order to increase power, we marked cells whose DNMT3A mutation status could not be determined by GoT as wild-type cells. For **Fig 2G**, we examined #122’s cells’ distribution across cell types.

### Sync-score

We defined the AVP signature to include genes with high correlation (> 0.6) to AVP across HSC, MPP and MEBEMP metacells, and the GATA1 signature to include those with correlation > 0.7 to GATA1. We filtered genes with mean relative expression > 2^-10 in these metacells, to preclude a small number of genes from dominating the signatures. We then selected HSC, MPP, MEBEMP-E and MEBEMP-L cells, scored them by the fraction of UMIs from the AVP and GATA1 signatures, and partitioned them into 20 bins of AVP signature expression and 20 bins of GATA1 signature expression, such that all AVP bins and all GATA1 bins had the same number of cells. The sync-score is then defined as the fraction of cells in GATA1 bin 13 and above (upper two quintiles of GATA1) that are in AVP bin 9 and above (upper three quintiles of AVP expression).To visualize the sync scores (**Fig 3I**), we normalized the 20×20 bin matrix to sum to 1, smoothed the obtained matrix by averaging cells using a running window of length 3, and taking log2.

### Differential gene expression with respect to age and CBC

Differential expression was performed separately for MPP / MEBEMP cells, and for CLPs. The MPP and CLP-M matrices that were used to detect variant gene modules were used here for differential expression. Differential expression was performed separately for males and females. Each gene’s expression value was correlated with max VAF, age and 20 CBC values using Spearman correlation, and the correlation was tested for significance. p-values were FDR-corrected (Benjamini-Hochberg) for each label separately. For max VAF we also performed a Mann-Whitney test comparing individuals with and without a detected mutation. Differential expression between sexes was done using Mann-Whitney test on the same expression matrices.

### Myeloid malignancies scRNA-seq initial processing

We processed additional 10x libraries using cellranger as described previously. As the myeloid malignancies data was multiplexed with other samples, including some that are not part of this study, we detected doublets using Vireo and Souporcell^27^, and assigned cells into individuals as we described above. All the malignancies data was sequenced with the Ultima platform and we corrected the UMI matrix by downsampling and resampling UMIs as described above. We then created a metacell model for each of 12 samples separately: 2 healthy individuals, 3 CMML patients, 2 MDS patients (one patient has two samples), 1 MDS / MPN patient, 1 myelofibrosis patient, and 2 AML patients. As before, we excluded cells with less than 500 UMIs, with more than 20% expression of mitochondrial genes, or with high expression of megakaryocyte genes. We used the same set of ignored genes as we use for the healthy PB metacell model. We set the target number of cell per metacell such that a metacell will have about 300K UMIs.

### Projection of disease data on the HSPC model

To project each individual’s metacells on the healthy reference, we correlated the query metacells with the reference’s metacells. The correlation was performed in log2 scale and after downsampling the query metacells to have 150K UMIs, using genes that were variable in the reference. Query metacells were then annotated using the most common cell type among the 5 atlas metacells they were most correlated with. Metacells that mapped to CD34- reference metacells were discarded for the following analyses. **Fig 4B** reports the mean correlation between each query metacell and its 5 most correlated reference metacells, and places each query metacell on the mean x and y coordinates of its 5 most highly correlated atlas metacells. To plot **Fig 4E**, we calculated the observed expression of each gene as the geometric mean expression across all metacells of an individuals that were annotated as MPP/MEBEMP-E/MEBEMP-L. We looked at the 5 most correlated atlas metacells for each one of these query metacells, and calculated for each gene its geometric mean to obtain the expected expression. Due to systematic differences between the query and atlas in some samples, we normalized the expected values by sorting genes based on their mean expression in the observed and expected, and subtracting from the expected value a rolling mean across 500 genes of the difference between the expected and observed. We defined differentially expressed (DE) genes as having an at least 2-fold difference between observed and expected.

**Figs 4C,D** are based on projection of single cells. Cells were correlated to metacells downsampled to 150K UMIs, and the raw UMI vectors were used for correlation. **Fig 4C** then shows the distribution of the annotation of the metacells that cells were projected onto. For **Fig 4D**, each query cell was classified to the bin (as in **Fig 2D**) that was most common among cells in the metacell to which it mapped.

### Karyotype analysis

To perform karyotype analysis, we subtracted from each metacell’s expression (log2 after normalizing to sum to 1) the geometric mean expression of the 5 atlas metacell it was most correlated with. For each metacell we then substracted the median difference value over all genes. We then partitioned each chromosome into bins with an equal number of genes such that all bins will have at least 40 genes, and computed the median difference across all genes in the bin. For this analysis, we only considered genes with an average expression of at least 2^-15.5 in either the pool of query metacells or the pool of the matched atlas metacells. This analysis provides metacell resolution karyotypes, as shown in **EDF 9**. To plot **Fig 4F** we calculated the median difference value for each gene across metacells that were projected to CD34+ cell types.

### Profiling signatures in disease cases

We separated AML-1 metacells into AML-1-1 and AML-1-2 by their karyotype as shown in **EDF 9**. We created lists of genes overexpressed in healthy NKTDPT, CLP, HSC and MEBEMP metacells. We scored each AML metacell by the geometric mean of all genes in each of the above gene lists, as well as in the LMNA signature genes. We set thresholds for a metacell to express a particular gene program by observing the distribution of the gene program’s expression across reference metacells in the relevant cell population (e.g., NKTDP metacells for the NKTDP gene program, see dashed lines in **EDF 10A**).

For the heatmap in **EDF 11A**, we selected genes from the above gene programs, as well as genes higher in AML-1 or in AML-2 compared to the reference across all cell types, and genes higher in AML-1-2 compared to AML-1-1 in all the following cell types: CLP-E, MPP and MEBEMP-E.

To discover de-novo gene programs in the AML samples (**EDF 10C**), we selected genes that are highly variable in each AML metacell model, calculated their correlation across metacells, clustered their correlation profiles, and filtered clusters with low mean correlation (less than 0.4), or genes with low mean correlation (less than 0.4) to their cluster. We manually added several genes of interest to the displayed matrices (**EDF 10C**).

## Data Availability

All data can be explored in: https://apps.tanaylab.com/MCV/blood_aging/

## Code Availability

Detailed code for the figures will be provided prior to publication.

The Metacell R package is available at https://github.com/tanaylab/metacell

## Contributions

Patient recruitment was performed by NF, LS, SB, and ST. Blood drawing and sample preparation for scRNAseq, including CD34 enrichment, were performed by NF. 10x scRNA library preparation and sequencing on the Illumina platform were performed by NF. scRNAseq data processing, including doublet calling, assignment of cells to individuals and metacell model construction were performed by NR. BM projections and comparisons were performed by NR. All compositional, differential gene expression and variant gene module analyses were performed by NR. Clinical data curation was performed by NF. Clinical data association analyses were performed by NR. MDS and AML patient –derived metacell projections and analyses were performed by NR. MCV construction was performed by AL. ARCH and genotyping deep targeted sequencing of all study participants were performed by NF. Amplicon validation of ARCH mutations was performed by NF. Sequencing data analysis and variant calling were performed by NC, NF and LS. DNA samples for the high-RDW-CH analysis were provided by SS and NA. ARCH sequencing of all high-RDW and control DNA samples was performed by NF and analyzed by NC, NF and LS. Replicate 10x library sequencing on the Ultima genomics platform was performed by ZS, DL, EM and GY. Biological and technical replicate analysis was performed by NR. GoT experiments were performed by SG, analyzed by ND and PC, supervised by DL and analyzed by NR. Gata3 deep targeted sequencing and Sanger validation were performed by SS and supervised by JT. AB, AL, and OBK contributed to data analysis and interpretation. AD provided sample preparation support. MK provided 10x guidance and technical support. NK contributed to funding applications. NF and NR wrote the manuscript. LS and AT designed and supervised all aspects of the study and wrote the manuscript.

## Corresponding authors

Correspondence to Liran Shlush and Amos Tanay

## Acknowledgements

This work was supported by the following grants: LLS and Rising Tide Foundation Grant ID: RTF6005-19, ISF-NSFC 2427/18, ISF-IPMP-Israel Precision Medicine Program 3165/19, ISF 1123/21, and the Ernest and Bonnie Beutler Research Program of Excellence in Genomic Medicine. LS is an incumbent of the Ruth and Louis Leland career development chair. This research was also supported by the Sagol Institute for Longevity Research, the Barry and Eleanore Reznik Family Cancer Research Fund, the Steven B. Rubenstein Research Fund for Leukemia and Other Blood Disorders, the Rising Tide Foundation and the Applebaum Foundation. The contribution of NR is part of a Ph.D. thesis research conducted at Tel Aviv University. NR was supported in part by a fellowship from the Edmond J. Safra Center for Bioinformatics, Tel Aviv University, and by the Planning and Budgeting Committee (PBC) fellowship for excellent PhD students in Data Sciences. NR was also supported by awards from the Herczeg Institute on Aging, and from the Tel Aviv University Healthy Longevity Research Center.

We thank all members of the Shlush and Tanay laboratories for their constructive comments.

## Supplementary Information

**Supplementary Table S1. – Basic participant demographics**. This table contains data on sex and age of all studied participants at sampling time, along with their allocated study IDs.

**Supplementary Table S2. – Longitudinal CBC’s**. This table contains median values for each CBC parameter for each sampled individual across 5 years prior to sampling date. WBC – white blood cells, RBC – red blood cells, MCV – mean corpuscular volume, MCH - mean corpuscular hemoglobin, MCHC - mean corpuscular hemoglobin concentration, RDW - RBC width distribution, MPV – mean platelet volume.

**Supplementary Table S3. – ARCH sequencing**. This table contains variant calling data on all ARCH positive individuals, including their Individual ID, specific gene involved, chromosome number, position, reference and alternate alleles, and the reported average variant allele frequency (VAF) from single (amplicon) or duplicate (MIP) sequencing instances.

**Supplementary Table S4. – HSC expression differences**. This table contains each gene’s mean expression across metacells in HSCs, MEBEMP-Es, and CLP-Ms, and the differences in mean expression between HSCs and the latter two cell types. These expression differences further classify genes as belonging to each of VIII groups as described in EDF 5B.

**Supplementary Table S5. – high RDW case-control longitudinal CBC’s**. This table contains all blood count instances of high RDW individuals and controls. WBC – white blood cells, RBC – red blood cells, MCV – mean corpuscular volume, MCH - mean corpuscular hemoglobin, MCHC - mean corpuscular hemoglobin concentration, RDW - RBC width distribution, MPV – mean platelet volume.

**Supplementary Table S6. – high RDW case-control ARCH sequencing**. This table contains variant calling data on all ARCH positive high-RDW cases and controls, including their Individual ID, specific gene involved, chromosome number, position, reference and alternate alleles, whether this alternation is considered a leukemic hotspot, and the reported average variant allele frequency (VAF) from single (amplicon) or duplicate (MIP) sequencing instances.

**Supplementary Table S7. – MEBEMP differential expression screen**. This table contains correlations between gene expression and different clinical labels across studied individuals. Correlations are between expression values normalized for distribution across the MPP-MEBEMP population, and the following labels:- maximal mutant VAF. The Spearman correlation between the maximal VAF and the gene expression is reported, as well as the p-value and BH corrected q-value for the correlation’s equality to 0 (two-sided test). These values are reported in columns starting with “vaf_cor”. The p-value and q-value are also reported for Mann-Whitney two-sided test, comparing the expression of the gene in individuals with and without mutations. These values are reported in columns starting with “vaf_mann_whitney”.- age. The Spearman correlation, p-value for correlation’s equality to 0 (two-sided) and BH-corrected q-value are reported for males and females separately. These are reported in the columns starting with “age”.- sex. The p-value and BH corrected q-value for a Mann-Whitney two-sided test comparing expression in males and females are reported. These are reported in the columns starting with “sex”.- CBCs. For each of 20 CBC values, and for males and females separately, three values are reported: the Spearman correlation between gene expression and CBC value, and p-value for that correlation being equal to 0, and the BH-corrected q-values. These values are reported in the remaining columns.Only genes whose expression levels exceeded a minimal threshold, and who displayed small technical variation, are listed.

**Supplementary Table S8. – CLP differential expression screen**. Similar to Supplementary Table 7, using CLP cell types.

**Supplementary Table S9. – LMNA gene correlations**. This table contains the Spearman correlation of each gene to *LMNA* across MEBEMP and CLP metacells (mc_cor columns), and across individual composition-controlled MEBEMP and CLP expression profiles (indiv_cor columns). The last column states whether the gene is part of the LMNA signature or not.

**Supplementary Table S10. – sync genes**. This table contains data for how the genes used to calculate the sync-scores were selected. It lists each gene’s correlation to AVP and GATA1 across the HSC-MPP-MEBEMP trajectory (avp_cor and gata1_cor), whether the gene is considered lateral and therefore excluded from the signature, mean expression across MEBEMP metacells, and whether the gene is part of the AVP and GATA1 gene signatures (is_avp_sig and is_gata1_sig).

**Supplementary Table S11. – Individual scores and signatures**. Key scores and signatures are listed for each individual: *LMNA* signature in MEBEMPs (lmna_mebemp_score), *LMNA* signature in CLPs (lmna_clp_score), sync-score, and the number of cells belonging to each cHSPC cell state (under columns whose names start with “num_”).

---

**EDF 1.**
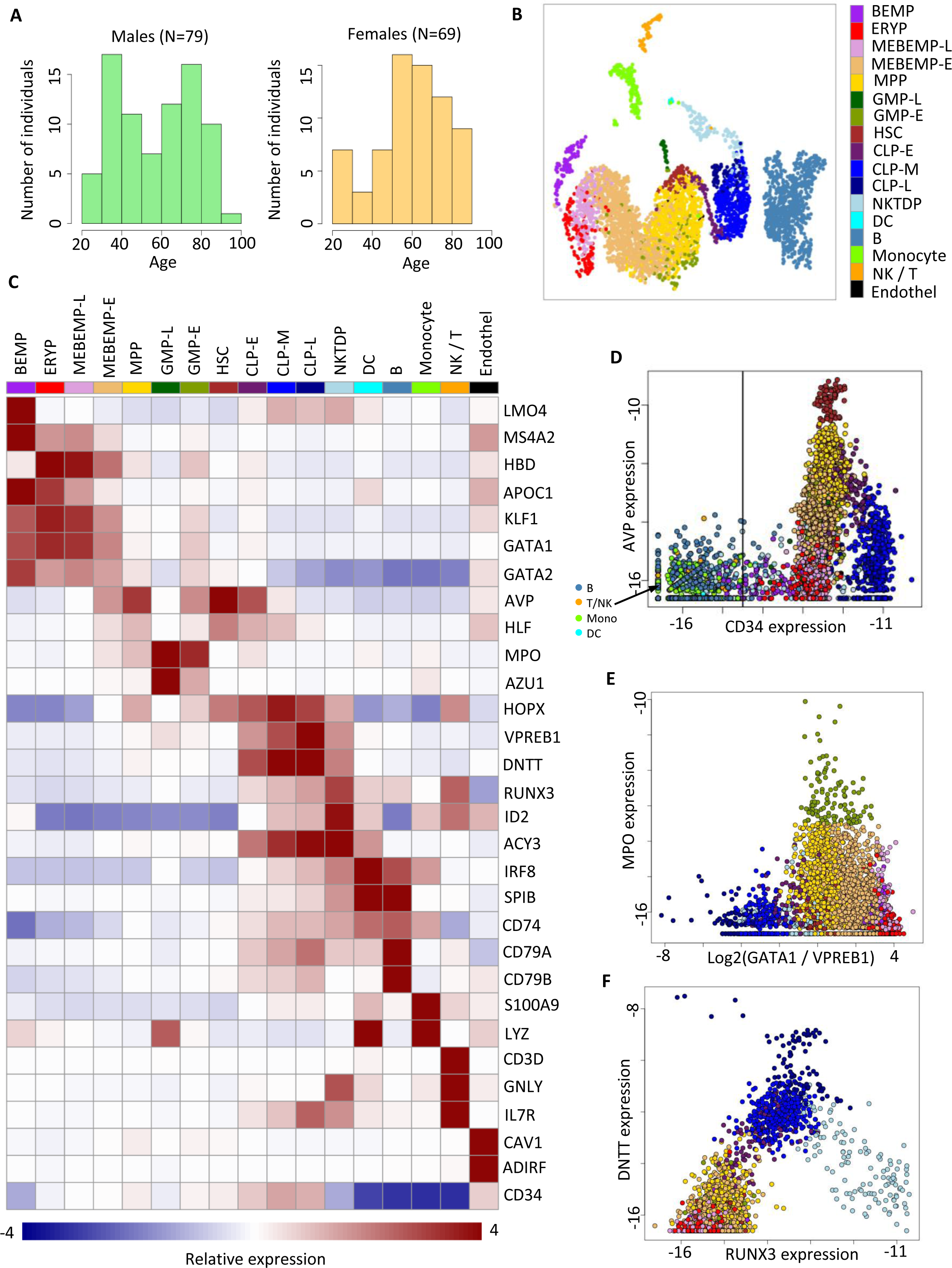
Cell state annotation, major markers and regulators of HSC differentiation and sub-population branching. **1A**– age distribution (decimals) of studied population by sex. **1B** – 2D UMAP projection of our metacell model prior to *CD34* metacell filtering. **1C** - relative expression heatmap of cell states (columns) and gene markers used for cell state annotation (rows). **1D** – Metacell filtration on CD34 levels. **1E** - expression plot of *MPO* and *GATA1/VPREB1* expression showing all 3 differentiation routes (GMPs, CLPs, MEBEMPs) from HSCs. **1F -** gene-gene expression plot of *DNTT* and *RUNX3*, showing early CLP differentiation and their bifurcation into late CLPs and NKTDPs. All gene expression values are obtained by normalizing gene expression to sum to 1 and taking log2.

**EDF 2.**
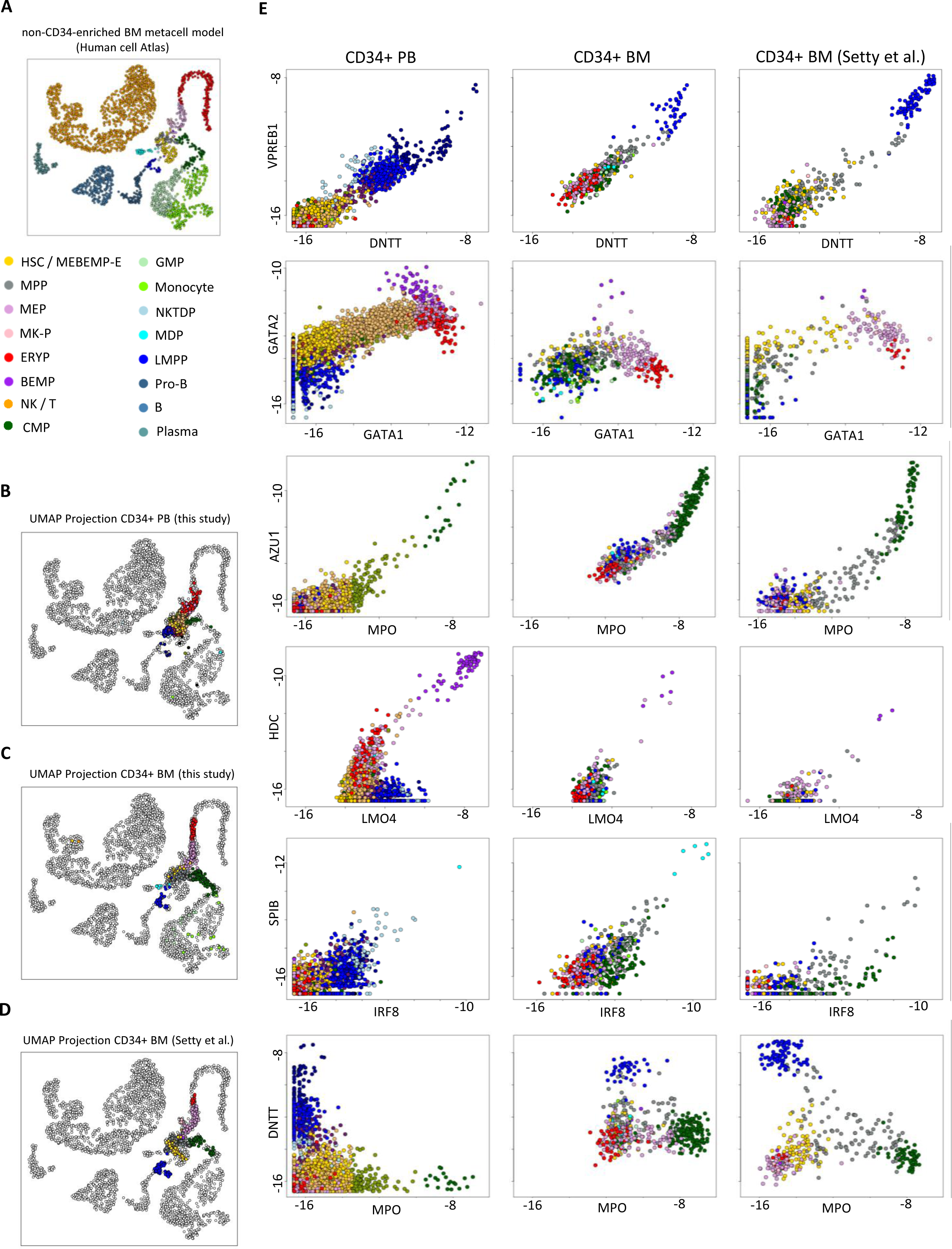
BM comparisons I. **2A** – 2D UMAP projection of a non-*CD34*-enriched BM metacell model from the Human Cell Atlas^16^, colored by a BM-specific cell state annotation. **2B** - projection of our PB *CD34*+ derived metacells on the non-*CD34* enriched BM metacell model. **2C** – projection of our BM *CD34*+ derived metacells on the non-*CD34* enriched BM metacell model. **2D -** projection of BM *CD34*+ derived metacells from Setty et al. on the non-*CD34* enriched BM metacell model. **2E** - gene-gene expression plots comparing PB *CD34*+ derived metacells with their BM *CD34*+ counterparts from our study and from Setty et al. for all differentiation trajectories. Panels (top to bottom) represent CLP differentiation, MEBEMP differentiation, GMP differentiation, BEMP differentiation, DC differentiation, and MPO/CLP/MEBEMP trifurcation from HSCs. PB / BM metacells are colored by PB / BM annotations, respectively.

**EDF 3.**
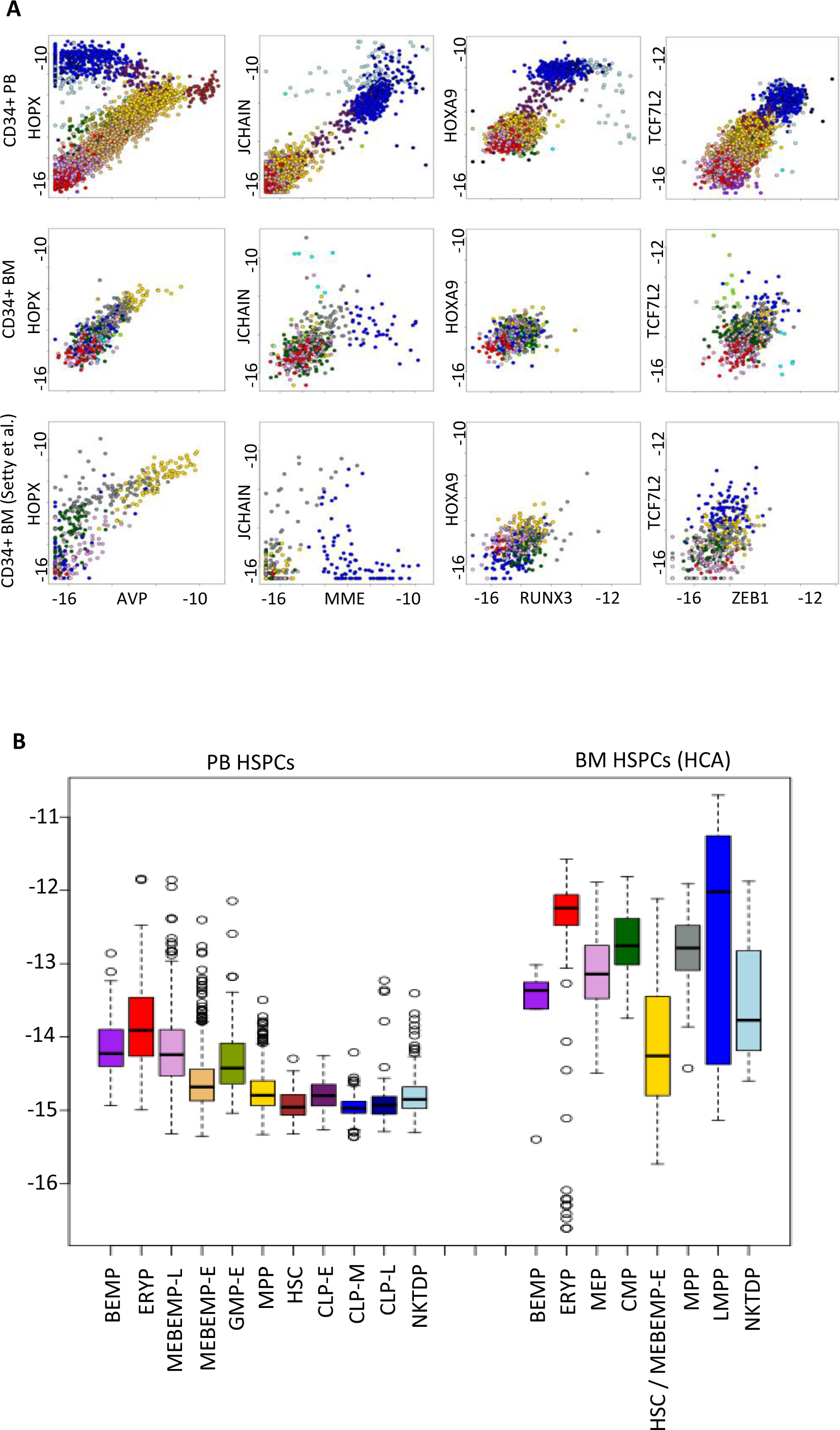
BM comparisons II – CLP differentiation and cell-cycling. **3A** – gene-gene expression plots comparing PB *CD34*+ derived metacells with their BM *CD34*+ counterparts from our study and from Setty et al. for markers and regulators of CLP differentiation and bifurcation, highlighting differences between PB and BM. **3B** – cell state specific comparison of S-phase signatures in circulating (left) vs. BM (HCA^16^, right) HSPCs.

**EDF 4.**
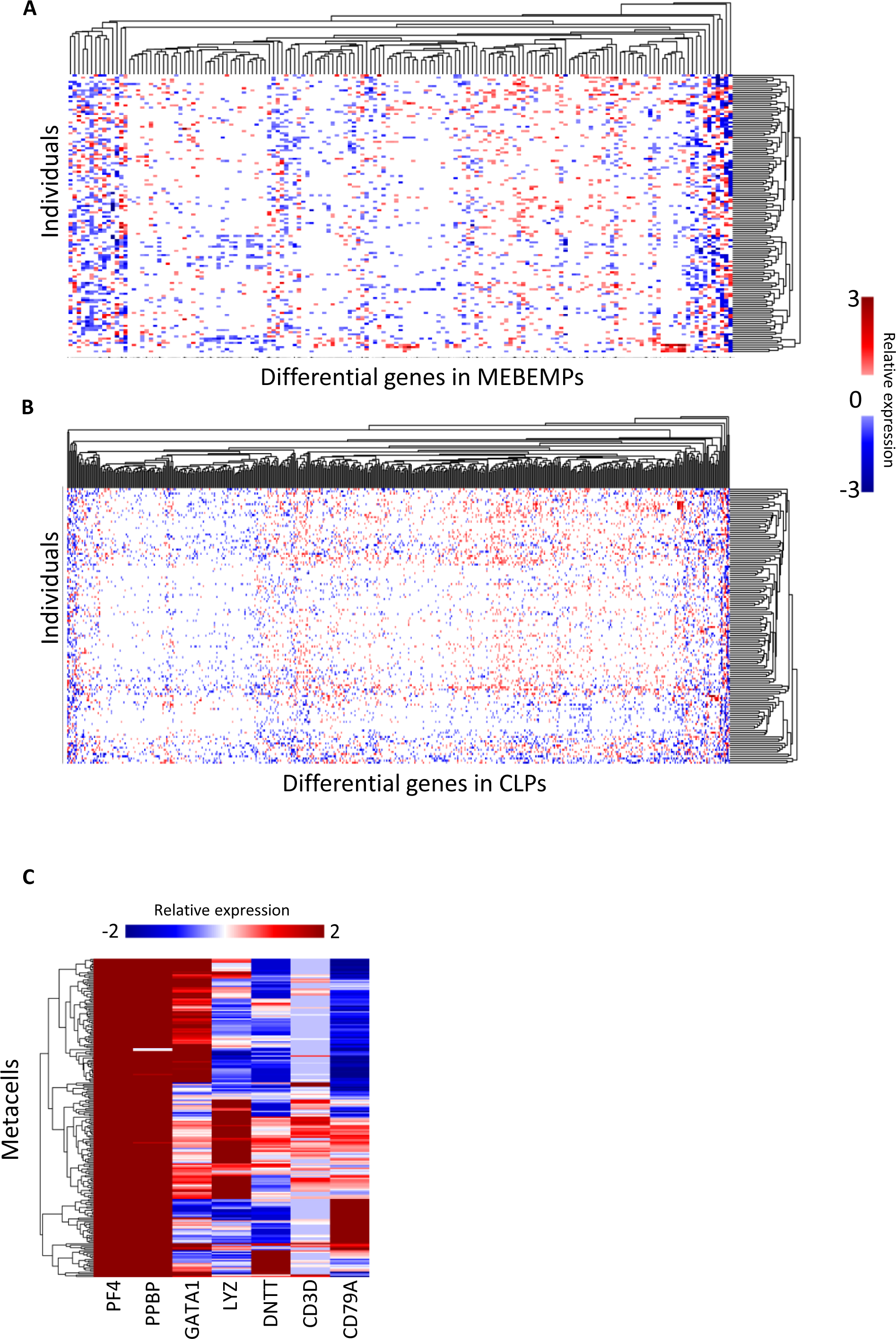
Individual-specific composition-controlled differential gene expression, megakaryocytic contamination. **4A,B** – Individual-specific differential gene expression after controlling for each individual’s distribution across the *CD34*+ PB manifold in MEBEMPs (top) and CLPs (bottom). **4C** - relative expression heatmap of the megakaryocytic markers *PF4* and *PPBP* and cell state specific markers, across metacells with high megakaryocytic signature. The figure shows an abnormally high doublet rate involving megakaryocytes. Cells contained in such metacells were accordingly excluded from the final metacell model.

**EDF 5.**
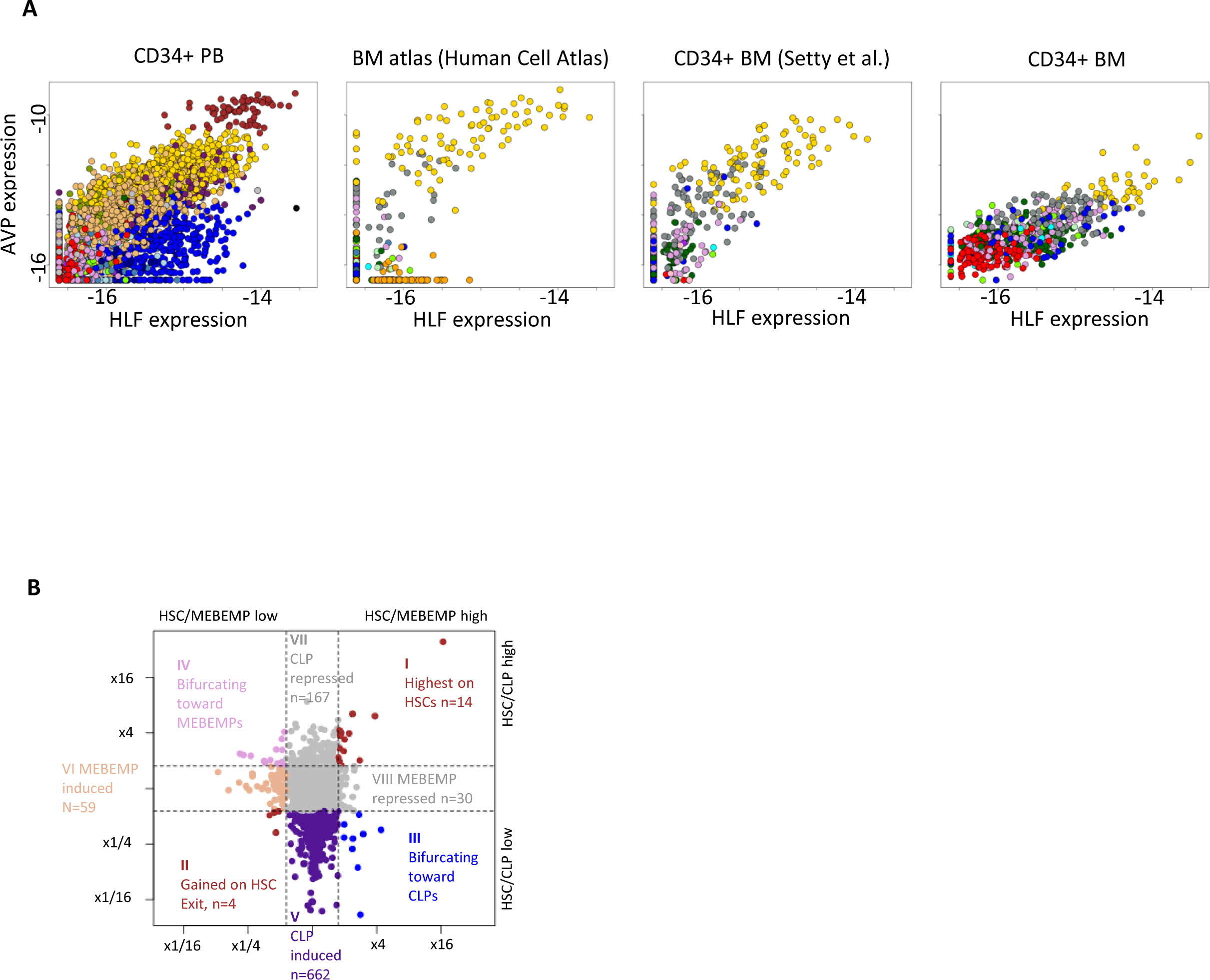
Circulating HSCs. **5A** - gene-gene expression plots comparing the PB high *AVP*/*HLF* HSC population (left) with that found in two BM metacell models ^16,17^ and in our CD34+ BM data. PB and BM metacells are colored by PB and BM annotations, respectively. **5B** – map of transcriptionally activated genes upon exit from the HSC state and differentiation toward lymphoid (CLP) and non-lymphoid (MEBEMP) fates. Dots represent genes. HSC/CLP and HSC/MEBEMP gene expression ratios are depicted on the y and x axis respectively. Class I genes are representative of the HSC state; Class II genes exhibit symmetric transcriptional activation upon exit from the HSC state towards CLP and MEBEMP fates, whereas Class III, IV, V, VI exhibit asymmetrical transcriptional activation upon exit from the HSC state towards CLP (class III, V) and MEBEMP (Class IV, VI) fates. n is the number of genes in each class.

**EDF 6.**
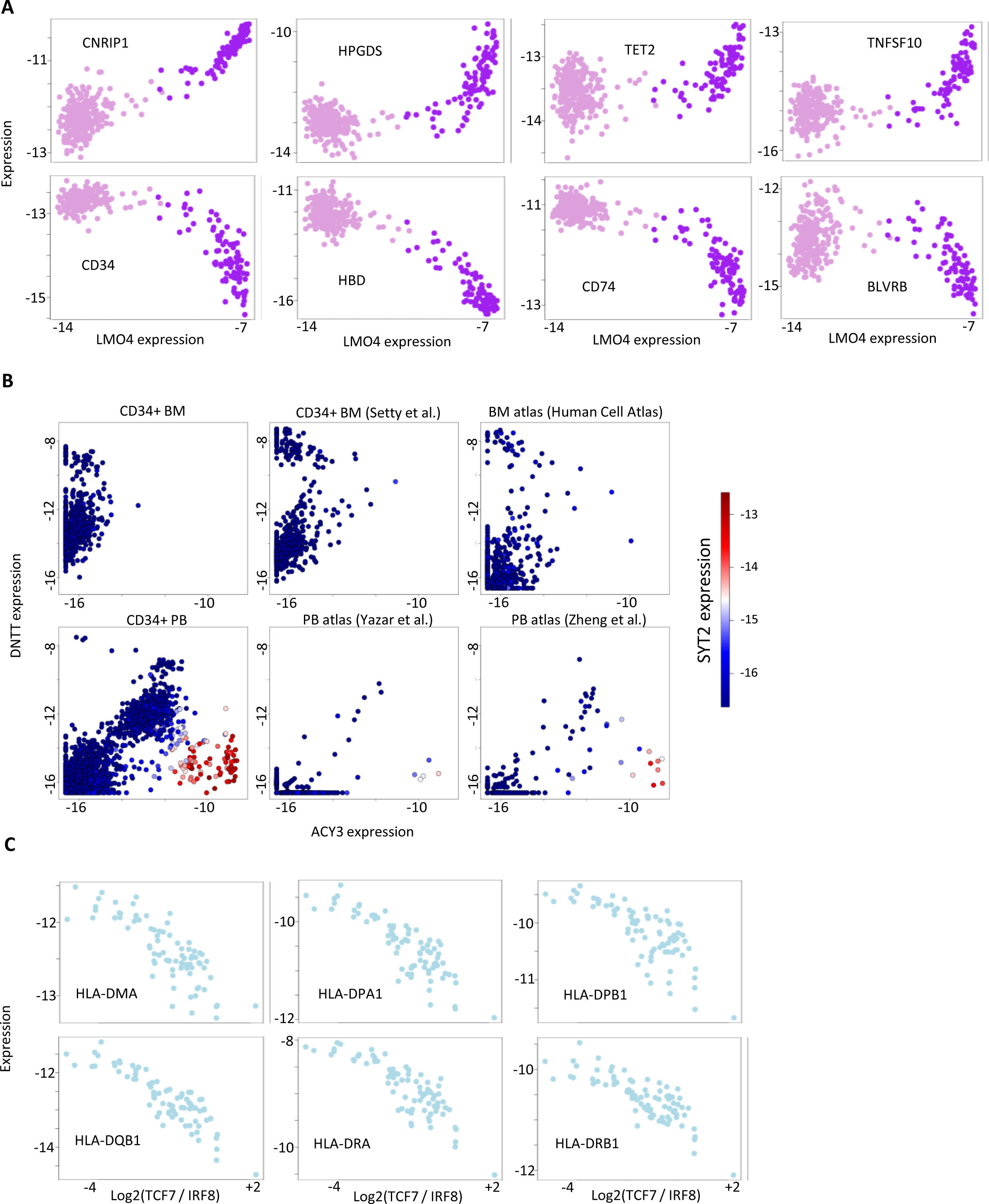
Factors involved in BEMP and NKTDP differentiation. **6A** - factors positively and negatively regulated in the early stages of BEMP specification. **6B** – gene-gene expression plots of *DNTT* and *ACY3* comparing *CD34*-enriched (Setty et al.^17^) and non-enriched (HCA^16^) BM to our *CD34*+ BM model (top), as well as non-enriched (Yazar et al.^28^) and partially enriched PB (Zheng et al.^36^) to our *CD34*+ PB model (bottom). Metacells are color-coded by *SYT2* expression. The *SYT2* high, *ACY3* high, *DNTT* intermediate population clearly seen in our data is completely lacking from the BM datasets. **6C** – anti-correlation of the DC *IRF8*-*MHC-II* coupled dynamics and the T cell regulator *TCF7*, involved in the bifurcation of the NKTDP cell state to its sub-populations.

**EDF 7.**
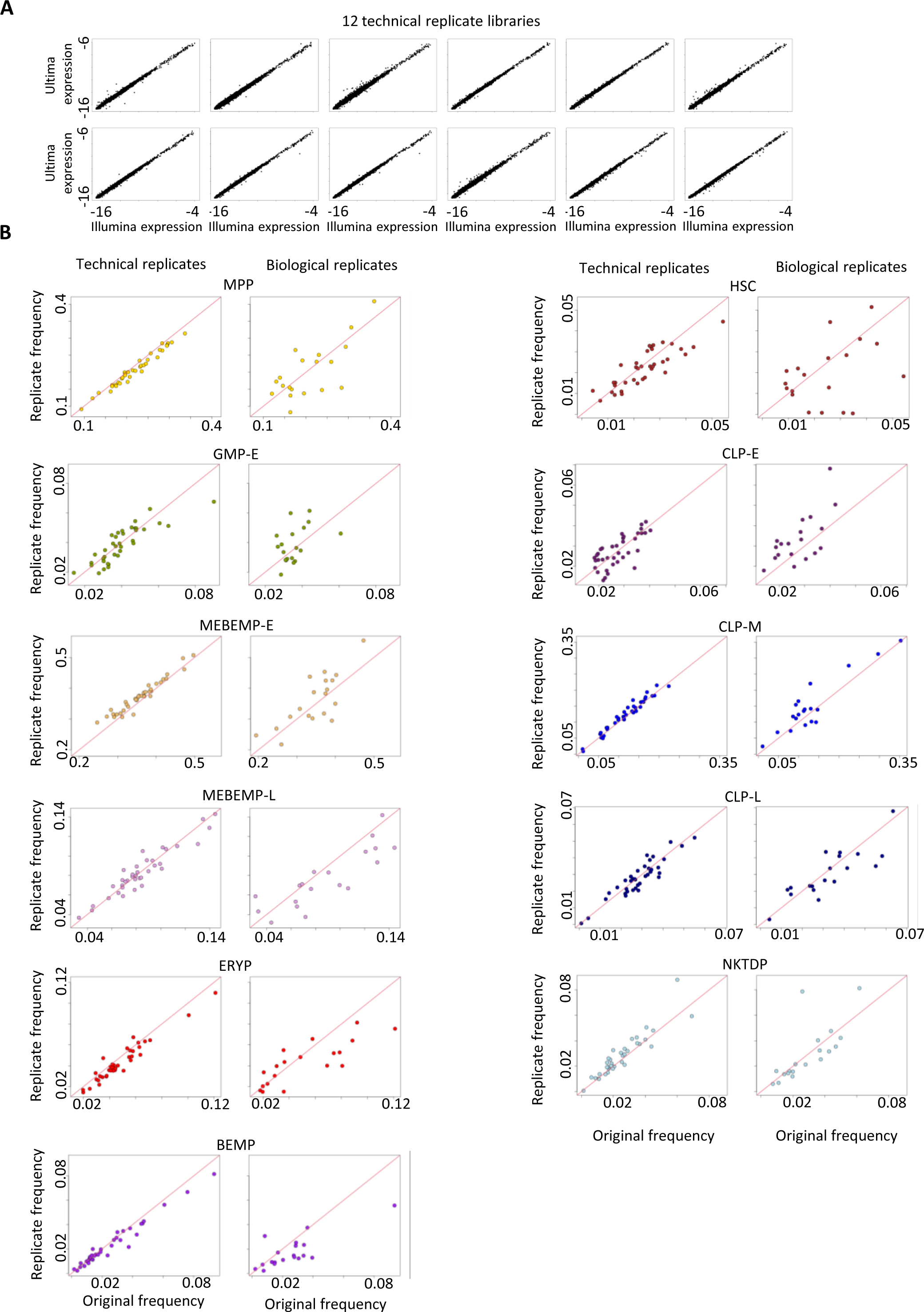
Stability of cell state compositions across technical and biological replicates. **7A** – comparison of Illumina and Ultima Genomics sequenced data. Each panel represents one library that was sequenced by both technologies. Points represent genes, and each gene’s expression level across all cells in the library as determined by Illumina (x) and by Ultima Genomics (y) is shown. **6B** – Cell state frequency comparisons between 39 technical & 20 biological replicates and their original samples. Each pair of panels represents one cell state, denoted on top. Panels on the left of each pair compare the cell state frequency in the original sample, sequenced by Illumina (x), to its technical replicate frequency, sequenced by Ultima Genomics (y). Panels on the right of each pair compare the cell state frequency in the original sample (x) to its biological replicate frequency (y). All biological replicates were sampled 1 year following original blood draw.

**EDF 8.**
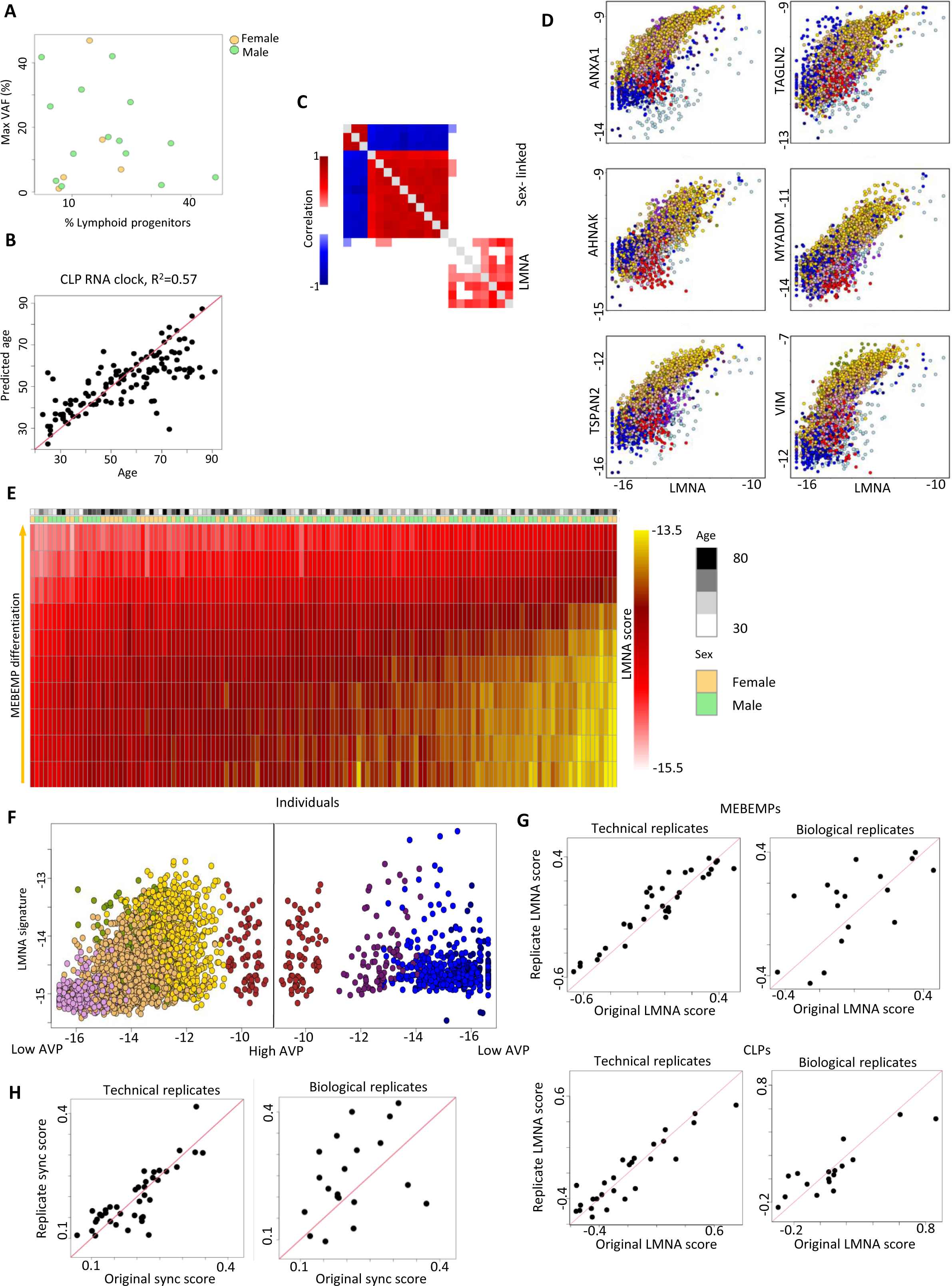
Composition-controlled transcriptional variation: the *LMNA* signature and sync score. **8A** – CLP frequency against maximal VAF (%) for individuals with CH. **8B** – True age (x) vs age predicted based on composition-controlled CLP expression (y). **8C** - gene-gene correlation heatmap, calculated over individual-level CLP gene expression controlled for CLP composition. **8D** – the *LMNA* signature – co-variation of *LMNA* expression with *ANXA1/2*, *AHNAK*, *MYADM, TSPAN2* and *VIM*. **8E** – heatmap of individual *LMNA* signature expression across the MEBEMP trajectory. Individual age and sex are color-coded on top. **8F** – *LMNA* signature expression in HSCs (denoted by high *AVP*) and throughout MPP / MEBEMP (left) and lymphoid (right) differentiation. **8G** - *LMNA* signature expression correlations between 39 technical & 20 biological replicates and their original samples. **8H** - sync score correlations between 39 technical & 20 biological replicates and their original samples. All biological replicates were sampled 1 year following original blood draw.

**EDF 9.**
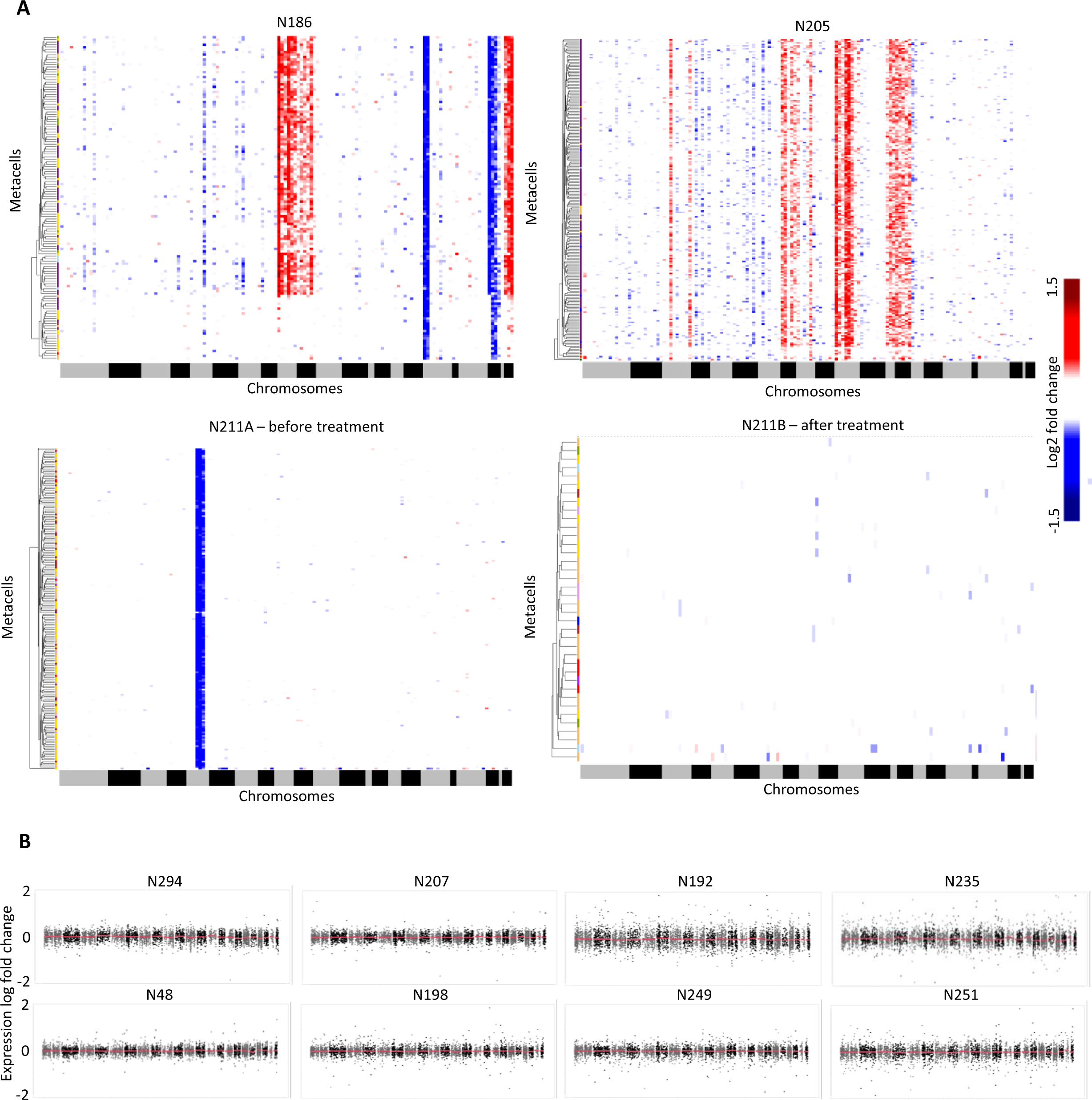
scRNAseq karyotyping. **9A** - scRNAseq karyotyping for 2 AML patients (N186, N205) and 1 MDS patient (N211) with isolated del5q (bottom), prior to and following treatment with lenalidomide. Metacell models were created for each MDS/AML patient and projected over our healthy reference map. Coupled reference and projected (patient) metacells were then used for calculating expression ratios over all expressed genes in all chromosomes. Heatmap of log2 expression fold-change (patient/reference) per metacell over all genes expressed in each chromosome is shown. **9B** – scRNAseq karyotyping for 2 healthy (N294, N207), 3 CMML (N192, N235, N48), 1 MDS (N249), 1 MDS/MPN (N251) and 1 Myelofibrosis (N198) patient. Karyotyping is performed as in 9A. Log2 fold-change expression (patient/healthy) for all expressed genes across all chromosomes is shown. Red lines represent the median of each chromosomal fold-change distribution.

**EDF 10.**
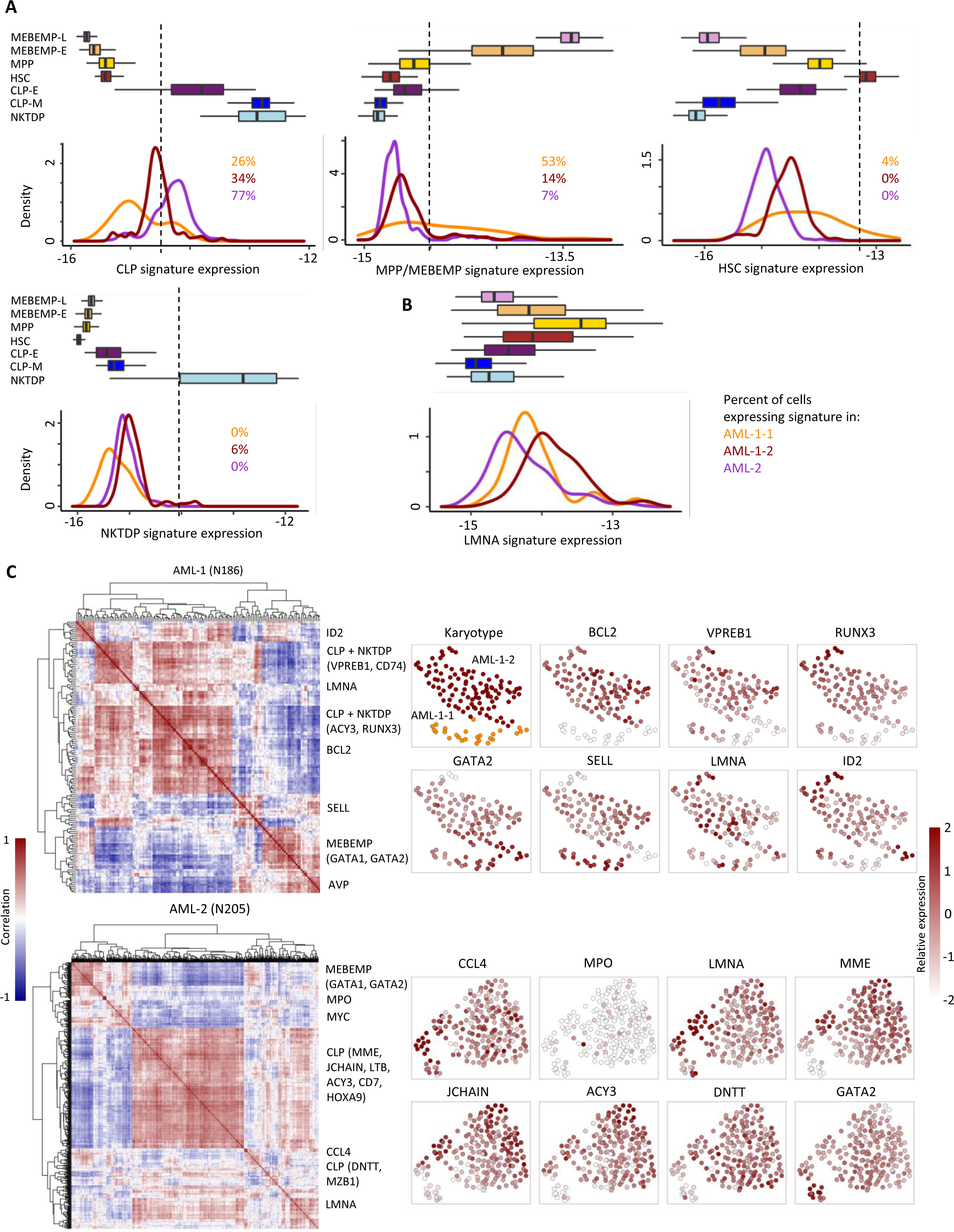
AML subpopulation analysis and characterization. **10A** - each of 4 panels refers to a different cell state gene signature as noted on the x-axis. Panel top - boxplots of gene module expression distributions for different cell states in our reference atlas. Panel bottom - Gene signature expression density plots for each of the AML subclones. Reference gene signature distributions (panel top) were used to identify subpopulations of AML cells with CLP, MEBEMP, HSC and NKTDP characteristics (panel bottom). Dashed lines represent the threshold for expressing a gene signature, and the fraction of cells expressing a signature per AML clone is listed. **10B** - same as 10A for *LMNA* expression across AML clones, showing high variability in *LMNA* expression across AML metacells. **10C** - left – correlation heatmap of differentially expressed gene signatures for AML-1 (N186, top) and AML-2 (N205, bottom). The malignant state is characterized by multiple novel gene signatures in addition to aberrant expression of “healthy” differentiation-related modules, right – UMAP projection of the metacell models of AML-1 (top) and AML-2 (bottom), colored by relative expression of differentially expressed genes. Overexpression of BCL2 in AML-1-2 compared to AML-1-1 can be seen.

**EDF 11.**
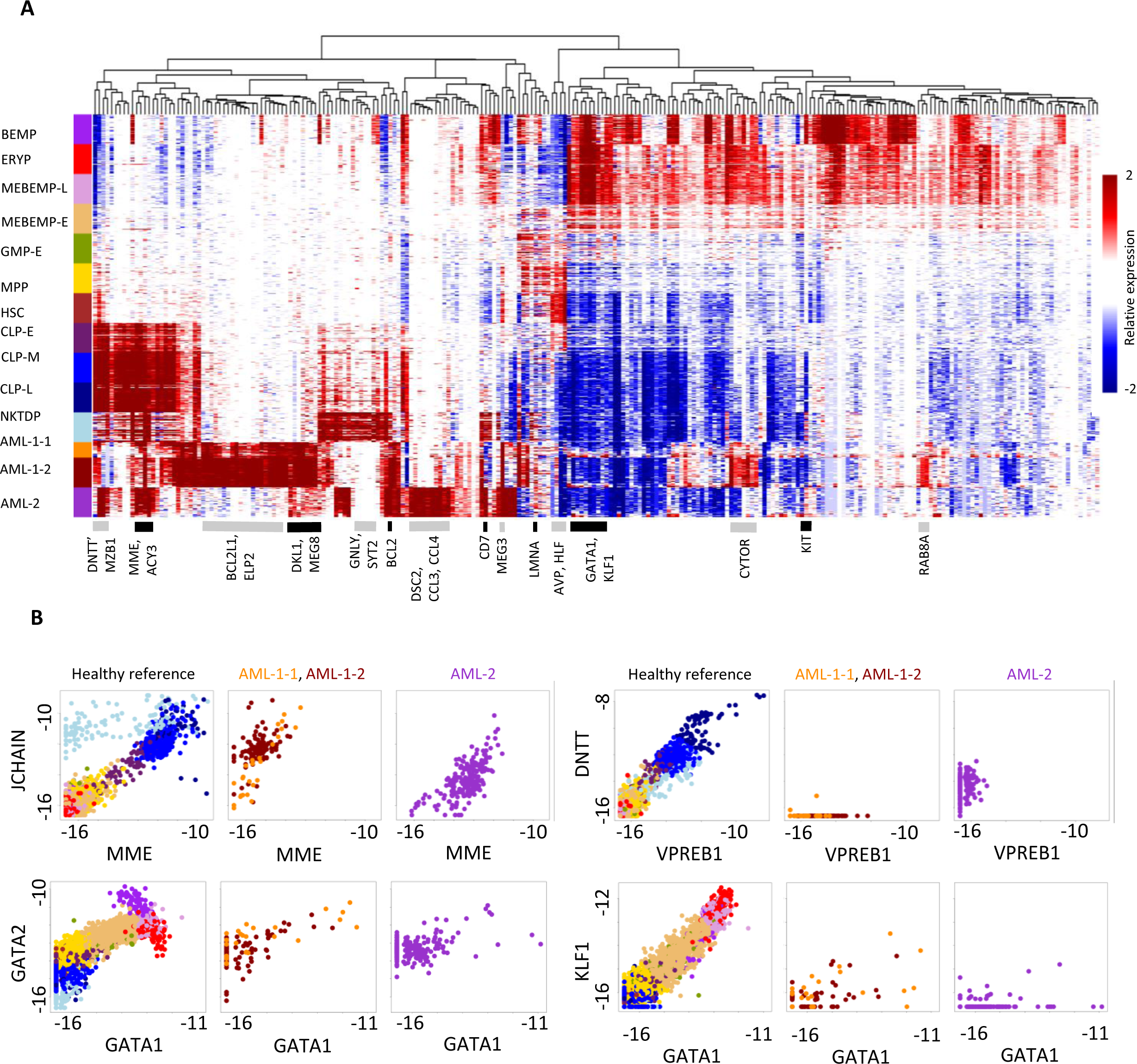
The transcriptional landscape of AML: aberrant expression of reference genes and multiple novel gene signatures. **11A** – Expression heatmap of several genes across reference cell states and AML subclones. The malignant state differs greatly from the healthy state both in the expression of reference genes and by multiple additional gene expression signatures. **11B** - panels compare gene expression of major differentiation regulators associated with lymphoid (top) and MPP / MEBEMP (bottom) differentiation, in reference (left), AML-1 (N186, middle, color coded by subclones) and AML-2 (N205, right) metacells.

